# Impaired phosphatidylethanolamine metabolism activates a reversible stress response that detects and resolves mutant mitochondrial precursors

**DOI:** 10.1101/2020.12.10.416495

**Authors:** Pingdewinde N. Sam, Elizabeth Calzada, Michelle Grace Acoba, Tian Zhao, Yasunori Watanabe, Anahita Nejatfard, Jonathan C. Trinidad, Timothy E. Shutt, Sonya E. Neal, Steven M. Claypool

## Abstract

Phosphatidylethanolamine made in mitochondria has long been recognized as an important precursor for phosphatidylcholine production that occurs in the endoplasmic reticulum (ER). Recently, the strict mitochondrial localization of the enzyme that makes PE in the mitochondrion, phosphatidylserine decarboxylase 1 (Psd1), was questioned. Since a dual localization of Psd1 to the ER would have far-reaching implications, we initiated our study to independently re-assess the subcellular distribution of Psd1. Our results support the unavoidable conclusion that the vast majority, if not all, of functional Psd1 resides in the mitochondrion. Through our efforts, we discovered that mutant forms of Psd1 that impair a self-processing step needed for it to become functional are dually localized to the ER when expressed in a PE-limiting environment. We conclude that severely impaired cellular PE metabolism provokes an ER-assisted adaptive response that is capable of identifying and resolving nonfunctional mitochondrial precursors.

## INTRODUCTION

Mitochondria are vital cellular components that generate energy, phospholipids, amino acids, reducing equivalents, and other essential metabolites. Nearly 99% of the mitochondrial proteome (~1000 proteins in yeast and ~1500 in mammals) is encoded in the nucleus, translated in the cytosol, and imported into one of four mitochondrial compartments—the outer membrane (OM), intermembrane space (IMS), inner membrane (IM), or matrix. Mitochondrial biogenesis is mediated by a series of dedicated import machineries in each compartment and is safeguarded by the collaborative efforts of an emerging network of factors including those that operate in other cellular compartments (Boos et al., 2019; Bykov et al., 2020; Hansen et al., 2018; Martensson et al., 2019; Matsumoto et al., 2019; Shpilka and Haynes, 2018; Wang and Chen, 2015; Weidberg and Amon, 2018; Wrobel et al., 2015). A small proportion of the nuclear-encoded mitochondrial proteome mediates and/or facilitates lipid biosynthetic or trafficking steps critical for the proper functioning of this organelle (Acoba et al., 2020; Sam et al., 2019). Mitochondrial lipid biosynthesis is dependent on the acquisition of phospholipid precursors provided by the endoplasmic reticulum (ER)/vacuole that fuel the generation of phosphatidic acid, phosphatidylglycerol, cardiolipin, and phosphatidylethanolamine (PE) (Elbaz-Alon et al., 2014; Honscher et al., 2014; Kawano et al., 2017; Kojima et al., 2016; Lu and Claypool, 2015). Mitochondrial PE is produced by phosphatidylserine decarboxylase-1 (Psd1), an enzyme that is embedded in the IM with its catalytic site in the IMS and which removes the carboxyl group from phosphatidylserine (PS) in either the IM or potentially the OM (Aaltonen et al., 2016; Tamura et al., 2012b). The newly-produced PE can then be exported from mitochondria and tri-methylated to phosphatidylcholine (PC) in the ER (Cui et al., 1993; Horvath et al., 2012; Samborski et al., 1990; Tamura et al., 2012b).

The Psd1 precursor contains a bipartite N-terminus that consists of a mitochondrial targeting signal (MTS) followed by a hydrophobic transmembrane domain (TM). Upon engaging the TIM23 translocase in the IM, the transmembrane domain located carboxy terminal to the MTS stops the translocation process and promotes lateral release of the Psd1 precursor into the IM (Horvath et al., 2012). Following two processing steps performed by matrix resident peptidases, membrane-anchored Psd1 performs a final self-processing event termed autocatalysis (Horvath et al., 2012; Onguka et al., 2015; Tamura et al., 2012b). Autocatalysis is mediated by an evolutionarily conserved catalytic triad that provides a single-use serine protease activity (Choi et al., 2015; Ogunbona et al., 2017; Watanabe et al., 2020). This self-processing event occurs at the conserved LGS motif and results in the formation of two subunits, α and β that remain non-covalently attached, and an N-terminal pyruvoyl prosthetic group on the α subunit that is essential for Psd1 activity (Li and Dowhan, 1988; Recsei and Snell, 1984). Interestingly, overexpression of mitochondrial precursor proteins with N-terminal bipartite signals, including Psd1, evokes a novel stress response called the mitochondrial Compromised Protein Import Response or mito-CPR (Weidberg and Amon, 2018).

Yeast lacking Psd1 have decreased respiratory capacity (Baker et al., 2016; Bottinger et al., 2012; Calzada et al., 2019) similar to that of *PISD-*deficient mammalian cells (Heden et al., 2019; Tasseva et al., 2013). Deletion of Psd1 also impairs autophagic capacity (Nebauer et al., 2007), decreases cell growth (Birner et al., 2001), reduces membrane fusion (Chan and McQuibban, 2012), and impairs OM protein biogenesis (Becker et al., 2013). In yeast, some of these defects can be completely or partially rescued through alternative PE biosynthetic pathways that reside in the ER, such as lyso-PE supplementation (Riekhof et al., 2007) or PE generation through the Kennedy pathway (Calzada et al., 2019; Riekhof and Voelker, 2006). Still, the limitations of these non-mitochondrial PE pathways to replace mitochondrial Psd1 are highlighted by the fact that murine whole body *pisd*^−/−^ mice die *in utero* and that rare pathogenic *PISD* mutations are associated with mitochondrial disease (Fullerton et al., 2007; Girisha et al., 2019; Peter et al., 2019; Selathurai et al., 2019; Steenbergen et al., 2005; Zhao et al., 2019).

PSDs are conserved from bacteria to humans and type I PSDs have been documented to localize to the mitochondrial IM in Arabidopsis (Rontein et al., 2003), *Saccharomyces cerevisiae* (Horvath et al., 2012; Tamura et al., 2012b), rats (Dennis and Kennedy, 1972; van Golde et al., 1974; Zborowski et al., 1983), hamster cells (Kuge et al., 1996; Voelker, 1985), and humans (Keckesova et al., 2017). Although there are Psd1 paralogs, classified as type II PSDs, that localize to endosomes in yeast (Psd2) (Gulshan et al., 2010) and to the tonoplast (PSD2) or ER (PSD3) in Arabidopsis (Nerlich et al., 2007), only the mitochondrial type I PSD is present in humans (PISD). The yeast mitochondrial Psd1 protein (Psd1) shares 48% identity with human PISD, while it only shares 24% identity with the non-mitochondrial Psd2 (Altschul et al., 1997; Altschul et al., 2005).

Tools to monitor phospholipid trafficking in real time are limited (Wills et al., 2018). As a surrogate, definitive information on the localization of enzymes in a sequential phospholipid metabolic pathway provides information on the trafficking steps that must have occurred. This paradigm has been extensively used to define steps involved in the PS to PE to PC pathway (Achleitner et al., 1995; Daum et al., 1986; Hovius et al., 1992; Kojima et al., 2016; Lahiri et al., 2014; Tamura et al., 2012a; Vance, 1991; Voelker, 1985). Further, it has served as the foundational premise for work leading to the molecular identification of the first mitochondria-ER contact site structure, ERMES (Kornmann et al., 2009; Nguyen et al., 2012), and was exploited to confirm the functional importance of contact sites between mitochondria and vacuoles (Elbaz-Alon et al., 2014; Honscher et al., 2014).

Recently, the foundation of this paradigm has been called into question. It was reported that a functionally significant fraction of Psd1 is N-glycosylated and thus targeted to the endomembrane system (Friedman et al., 2018). Moreover, the relative abundance of glycosylated Psd1 was reported to be sensitive to metabolic growth conditions, being most abundant when yeast cells are grown on fermentable carbon sources, like dextrose. Combined with studies that employed Psd1 chimeras targeted to either the ER or IM, the metabolically-sensitive distribution of Psd1 was taken as evidence that Psd1 produces two distinct subcellular pools of PE that have non-redundant functions (Friedman et al., 2018).

Given how a dual localization of Psd1 would impact the interpretation of numerous studies relying on its strict mitochondrial localization, here, using yeast and mammalian models, we re-examined its subcellular localization and determined that wild type (WT) Psd1 co-fractionates and co-localizes exclusively with mitochondria. Multiple Psd1 chimeras targeted to the IM via sequence elements provided by other topologically similar IM proteins functionally replace WT Psd1. Interestingly, we confirmed that a significant proportion of an autocatalytic mutant of Psd1, which is enzymatically non-functional, is N-glycosylated and thus ER-targeted (Friedman et al., 2018). Additional mutations that impair autocatalysis are also partially directed to the ER. In contrast to the mitochondrial targeted autocatalytic mutant Psd1, the glycosylated mutant Psd1 present in the ER is ubiquitinated and rapidly degraded, likely by the proteasome. The accumulation of glycosylated non-functional Psd1 in the ER only occurs when cellular PE metabolism is significantly disrupted and can be prevented by supplements that stimulate different lipid biosynthetic pathways that reside in the ER and that do so via distinct mechanisms. In addition to re-establishing the strict mitochondrial localization of functional Psd1, our work supports that Psd1 quality assurance is in part monitored during its import and enforced with the assistance of the ER and ubiquitin-proteasome system. We conclude that severely impaired cellular PE metabolism provokes an adaptive response that is capable of identifying and resolving nonfunctional mitochondrial precursors.

## RESULTS

### Functional Psd1 is a mitochondrial resident

Given the recent suggestion that Psd1 has dual residences (Friedman et al., 2018), we decided to thoroughly re-evaluate its subcellular localization using multiple distinct yeast strain backgrounds. First, we compared the specificity of two in-house generated Psd1-specific antibodies raised in two different rabbits (4077 and 4078) with similar sensitivities (Figure 1A). Notably, although each antiserum recognized the Psd1 β subunit in mitochondria, they each cross-reacted with several closely-migrating background bands that were importantly not the same. Thus, each antiserum specifically recognizes Psd1 β but also reacts with different non-Psd1 proteins.

**Figure 1.**
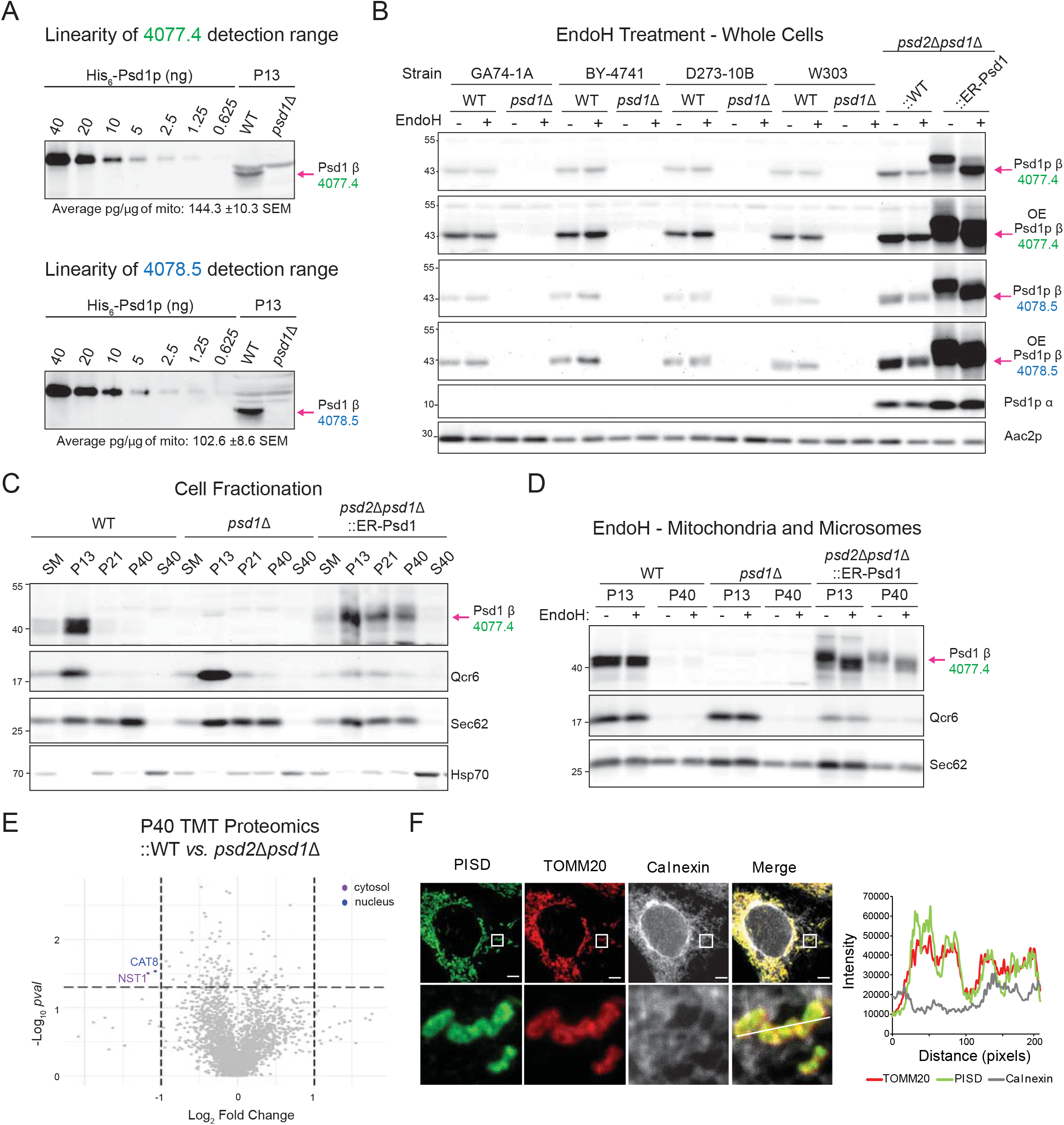
Endogenous and overexpressed Psd1 is localized to mitochondria. (A) Psd1 detection range was determined for antisera raised in two rabbits 4077 (*green*) and 4078 (*blue*) using a standard curve of recombinant Psd1. Bleeds number 4 and 5 were analyzed, respectively. The amount of Psd1 detected in WT mitochondria was calculated (n=3). (B) Cell extracts derived from pairs of WT and *psd1*Δ yeast of the indicated backgrounds grown at 30°C in synthetic complete dextrose (SCD) were treated with EndoH as listed and analyzed by immunoblot using the designated anti-Psd1 antisera; Aac2 served as loading control. *psd2*Δ*psd1*Δ yeast stably transformed with WT (∷WT) or ER-targeted Psd1 (∷ER-Psd1) acted as overexpression and glycosylation controls, respectively. OE indicates overexposed blots (n=3). (C) Following growth in SCD medium to late log phase, fractions were collected from the indicated yeast strains by differential gravity centrifugation. Equal protein amounts from each collected fraction were resolved by SDS-PAGE and immunoblotted for Psd1 and mitochondrial (Qcr6), ER (Sec62), and cytosolic (Hsp70) markers. SM, starting material, P13, pellet of 13,000x*g*; P21, pellet of 21,000x*g*; P40, pellet of 40,000x*g*; and S40, supernatant of 40,000x*g*. (n=3). (D) Mitochondria (P13) and ER (P40) fractions were mock or EndoH treated prior to immunoblot analysis (n=3). (E) Tandem mass tag (TMT) comparison of ER (P40) proteomes from *psd2*Δ*psd1*Δ (n = 3 preps) and *psd2*Δ*psd1*Δ∷WT yeast (n=4 preps). (F) Representative images of HEK293 cells overexpressing FLAG-tagged WT PISD obtained via immunofluorescence to visualize PISD (anti-FLAG; *green*), mitochondria (anti-TOMM20; *red*) and ER (anti-calnexin; *grey*). Bottom panels are a magnification of the white-boxed areas shown in the upper panel. Intensity profile for PISD (*green*), mitochondria (*red*) and ER (*grey*) along the pixels indicated by a solid white line.

Next, we tested the N-glycosylation status of endogenous Psd1 using endoglycosidase H (EndoH) in cell extracts derived from four different yeast strain backgrounds― GA74-1A, BY-4741, D273-10B, and W303― grown in synthetic complete dextrose (SCD) medium (Figure 1B). Neither the same antiserum (4077) as used by (Friedman et al., 2018) nor our second polyclonal Psd1 antibody (4078), detected an EndoH-sensitive form of Psd1 in any of the tested WT backgrounds. As a glycosylation control, we analyzed a *psd2*Δ*psd1*Δ strain expressing an ER-targeted Psd1 chimera (designated ∷ER-Psd1) which as expected (Onguka et al., 2015), was glycosidase-sensitive. Notably, a glycosylated form of Psd1 was also not detected in *psd2*Δ*psd1*Δ yeast expressing WT Psd1 (noted as ∷WT) at higher than endogenous levels, or in WT and *psd1*Δ strains employed in the Friedman study (Figure S1; (Friedman et al., 2018)).

If Psd1 is glycosylated, it should accumulate in membrane fractions that are lighter than mitochondria. However, endogenous Psd1 in GA74-1A yeast grown in rich dextrose media (YPD) did not co-fractionate with the microsomal fraction (P40) following differential centrifugation, in contrast to ER-Psd1 (Figure 1C). Consistently, we did not detect an EndoH-sensitive form of WT Psd1 in either the P13 (mitochondria and ER) or P40 fractions (Figure 1D). In contrast, ER-Psd1, which co-fractionated with ER-associated mitochondria (P13) and microsome-enriched membranes (P40), was sensitive to the glycosidase. Further, a tandem mass tag (TMT) proteomics comparison of P40 fractions from ∷WT and *psd2*Δ*psd1*Δ failed to detect Psd1-derived tryptic peptides in the microsome fractions from the yeast overexpressing WT Psd1 (Figure 1E). Our failure to detect Psd1, glycosylated or otherwise, in microsomes is consistent with the lack of significant Psd1 activity in this compartment when the non-mitochondrial Psd2 is missing (Onguka et al., 2015; Trotter and Voelker, 1995).

Finally, to extend our analysis to human cells, the localization of human PISD was determined by super-resolution microscopy in HEK293 cells over-expressing PISD harboring a C-terminal FLAG tag (Figure 1F). The anti-FLAG signal for PISD (*green*) overlapped with the mitochondrial marker, TOMM20 (*red*), but not with the ER marker, calnexin (Figure 1F). Immunofluorescence signal intensity plots of super-resolution images quantitatively showed that WT human PISD sharply overlapped with a mitochondrial marker. Collectively, our results support the unavoidable conclusion that the vast majority, if not all, functional Psd1 resides in the mitochondrion.

### Psd1 targeted to, and embedded in, the IM via non-Psd1 signals are functional

Friedman *et al.* used a chimeric strategy in which Psd1 was either directed to the ER or the IM, the latter by virtue of replacing the MTS and TM of Psd1 with the equivalent information from Mic60, and concluded that Psd1 targeted to either the mitochondria or the ER produces two distinct subcellular pools of PE that have non-redundant functions (Friedman et al., 2018). Specifically, when chimeric Psd1 is forced to exclusively localize to the IM, it is fully capable of supporting respiratory growth that requires mitochondrial energy production; however, it is unable to promote robust growth on dextrose that does not depend on energy from oxidative phosphorylation (Friedman et al., 2018). Given the significance of the reported growth phenotype with respect to Psd1 function and the underlying lipid trafficking steps needed to support its activity, we decided to adopt and expand upon this chimeric strategy. To this end, we designed a series of constructs, each containing a C-terminal 3XFLAG tag to monitor autocatalysis, in which the N-terminal MTS and TM domains of Psd1 were replaced by equivalent components from other inner membrane proteins with the same topology (Tim50, Yme1, Tim54, Mic60, and Ccp1) (Figure 2A). Each chimera, similar to previously characterized OM-Psd1 (Calzada et al., 2019) and ER-Psd1 (Onguka et al., 2015), produced Psd1 β subunits that predictably varied in molecular weight and α subunits that were the same size, indicating that they each retained self-processing activity (Figure 2B). Consistent with their autocatalytic status, each chimera rescued the *psd2*Δ*psd1*Δ ethanolamine auxotrophy (Figure 2C) and generated mitochondrial PE levels that were indistinguishable from ∷WT, with only one exception (Figures 2D and E). The elevated mitochondrial PE in ∷54-Psd1 mitochondria is highly reminiscent of what we previously observed when Psd1 is re-directed to the OM (OM-Psd1; (Calzada et al., 2019)). As such, we determined the subcellular and submitochondrial localization of each chimera in comparison to WT Psd1. The β and α subunits of each chimera co-fractionated with the mitochondrial inner membrane marker, Qcr6, and not the endosomal marker, Sec62 (Figure S2A). Using a well-established protease protection assay in isolated mitochondria, we determined that like WT Psd1, each chimera, with the possible exceptions of 54-Psd1 and Ccp1-Psd1, was protected from protease in intact mitochondria (Mito), similar to IM (Tim54) and matrix (Abf2) proteins, and unlike proteins in the OM (Tom70) (Figure S2B). Of note, a modest fraction of the 54-Psd1 chimera was degraded in intact mitochondria treated with protease, which could indicate that a proportion of this chimera is stuck in the OM and thus explain its elevated mitochondrial PE levels (Figure 2E).

**Figure 2.**
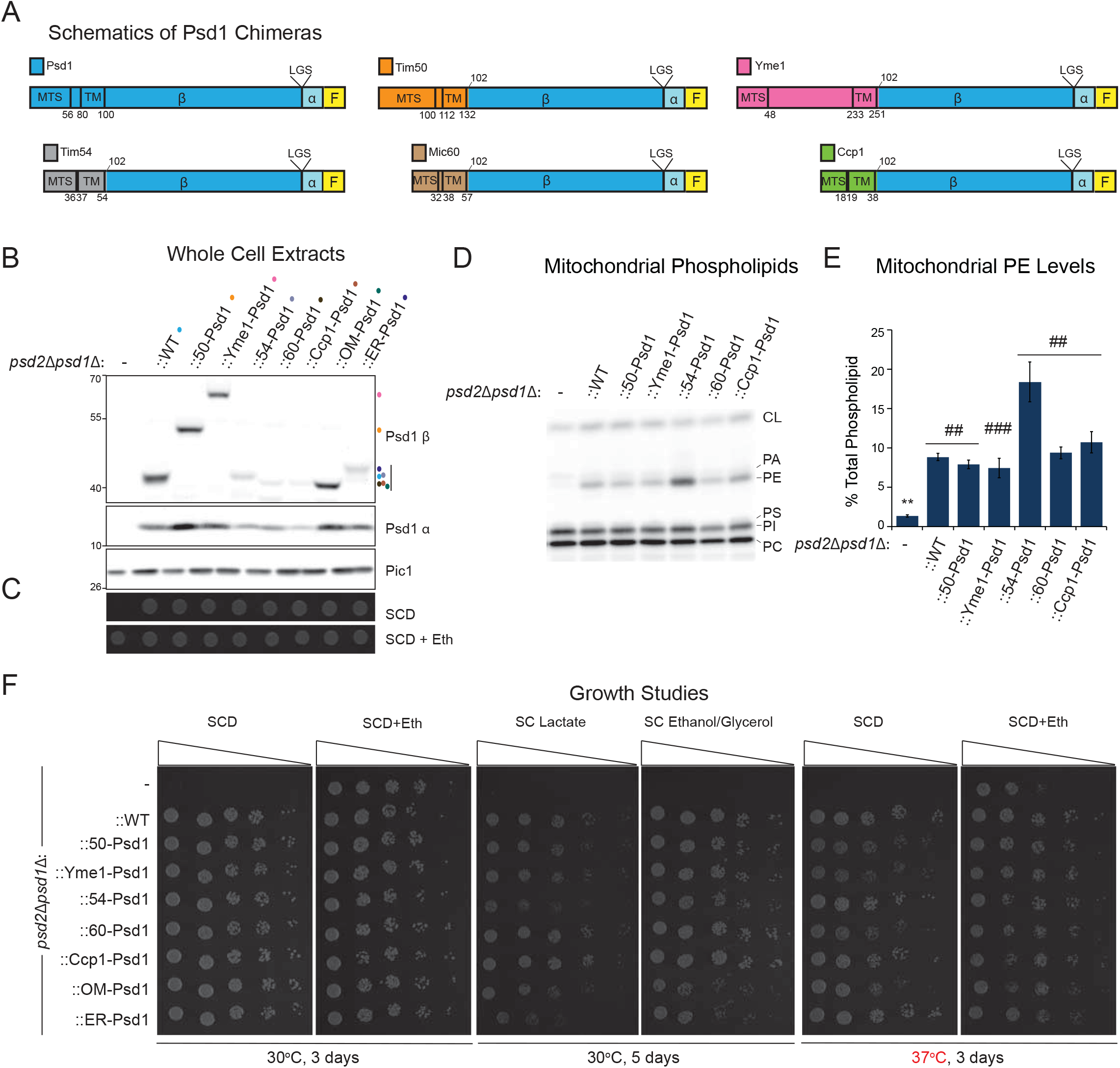
Re-localized Psd1 constructs are stable and functional. (A) Schematic of chimeric constructs. MTS (mitochondrial targeting signal) and TM (transmembrane domain) of residues are indicated. Psd subunits β, α and LGST motif are shown. All constructs have a 3XFLAG tag at the C-terminus. (B) The indicated strains were pre-cultured at 30°C in YPD, and after isolation of whole-cell extracts, the α and β subunits of Psd1 were analyzed by immunoblotting. Pic1 served as a loading control (n=3). (C) The indicated strains pre-cultured at 30°C in YPD were spotted onto SCD with (+) or without (-) 2 mM ethanolamine and incubated at 30°C for 3 days (n=3). (D) Mitochondrial phospholipids from the indicated strains were labeled overnight in rich lactate medium spiked with 32Pi and separated by TLC. The migration of phosphatidylcholine (PC), phosphatidylinositol (PI), PS, phosphatidylglycerol (PG), PE, phosphatidic acid (PA), and CL is indicated (n=6). (E) The relative abundance of PE was determined for each strain (mean ± SPEM for n=6). Statistical differences (2 symbols P ≤ 0.01; 3 symbols P ≤ 0.001 compared to ∷WT (*) or *psd2*Δ*psd1*Δ (#) were calculated by unpaired Student's t-test (for both ∷Yme1 comparisons) or Mann-Whitney Rank Sum Test (all the rest). (F) Serial dilutions of the indicated strains were spotted onto SCD with or without 2 mM ethanolamine, synthetic complete lactate (SC Lactate), or synthetic complete ethanol-glycerol (SCEG) plates and incubated at 30°C or 37°C for the indicated duration (n=3).

Finally, we determined the growth phenotypes of *psd2*Δ*psd1*Δ yeast expressing the assorted chimeras in comparison to WT Psd1, OM-Psd1, and ER-Psd1 on synthetic media containing dextrose with or without ethanolamine, lactate, or ethanol glycerol (Figure 2F). Each chimera, including Mic60-Psd1, grew similar to each other and to WT Psd1 in SCD with or without ethanolamine at 30°, but some chimeras (50-Psd1, Yme1-Psd1, 60-Psd1, and OM-Psd1) displayed a slight temperature-sensitive phenotype at 37°C (Figure 2F). With the sole exception of 54-Psd1, each chimera rescued the severe respiratory growth phenotype of *psd2*Δ*psd1*Δ yeast similar to WT Psd1. In contrast, ∷54-Psd1 growth was delayed compared to WT Psd1 (and the other chimeras) in respiratory conditions (SC Lactate and SCEG) (Figure 2F). This growth phenotype correlates with the previous observation that strains with elevated mitochondrial PE levels (such as OM-Psd1 and ER-Psd1) have diminished respiratory capacity (Calzada et al., 2019). Our combined results indicate that Psd1 targeted to and embedded in the inner membrane via non-Psd1 elements functions for the most part the same as WT Psd1. Based on the absence of any overt phenotype, we conclude that Psd1 contained within mitochondria is capable of supporting all activities, mitochondrial and non-mitochondrial, that depend on the PE that it produces.

### Glycosylation of mutant Psd1 requires severe cellular membrane dyshomeostasis

Psd1 autocatalysis is necessary for PE synthesis and amino acids required for this process have been functionally characterized (Choi et al., 2015; Ogunbona et al., 2017) including residues at the site of cleavage (LGS motif) and catalytic residues in its vicinity (Watanabe et al., 2020). With the goal of determining whether or not the putative glycosylated Psd1 and assorted IM- and ER-directed chimeras were capable of autocatalysis, non-functional, S463A mutant forms of Psd1, unable to perform autocatalysis, were used as immunoblotting controls and yielded Psd1 products that indeed migrated slower than their fully processed counterparts (Friedman et al., 2018). Notably, the relative amount of the glycosylated non-functional Psd1-S463A was significantly higher than that of the WT protein.

Therefore, to confirm and potentially extend this previous observation, we analyzed the glycosylation status of Psd1 mutants for each amino acid of the catalytic triad― S463A, H345A, and D120A. Consistent with (Friedman et al., 2018), we observed both EndoH-resistant (Mito β-α; *blue arrow*) and EndoH-sensitive (ER β-α; *purple brackets*) unprocessed forms for all three catalytic triad mutants (Figure 3A). These higher molecular weight protein bands were not detected in ∷WT where only mature Psd1 β (Mito β; *pink arrow*) and the released Psd1 α were detected.

**Figure 3.**
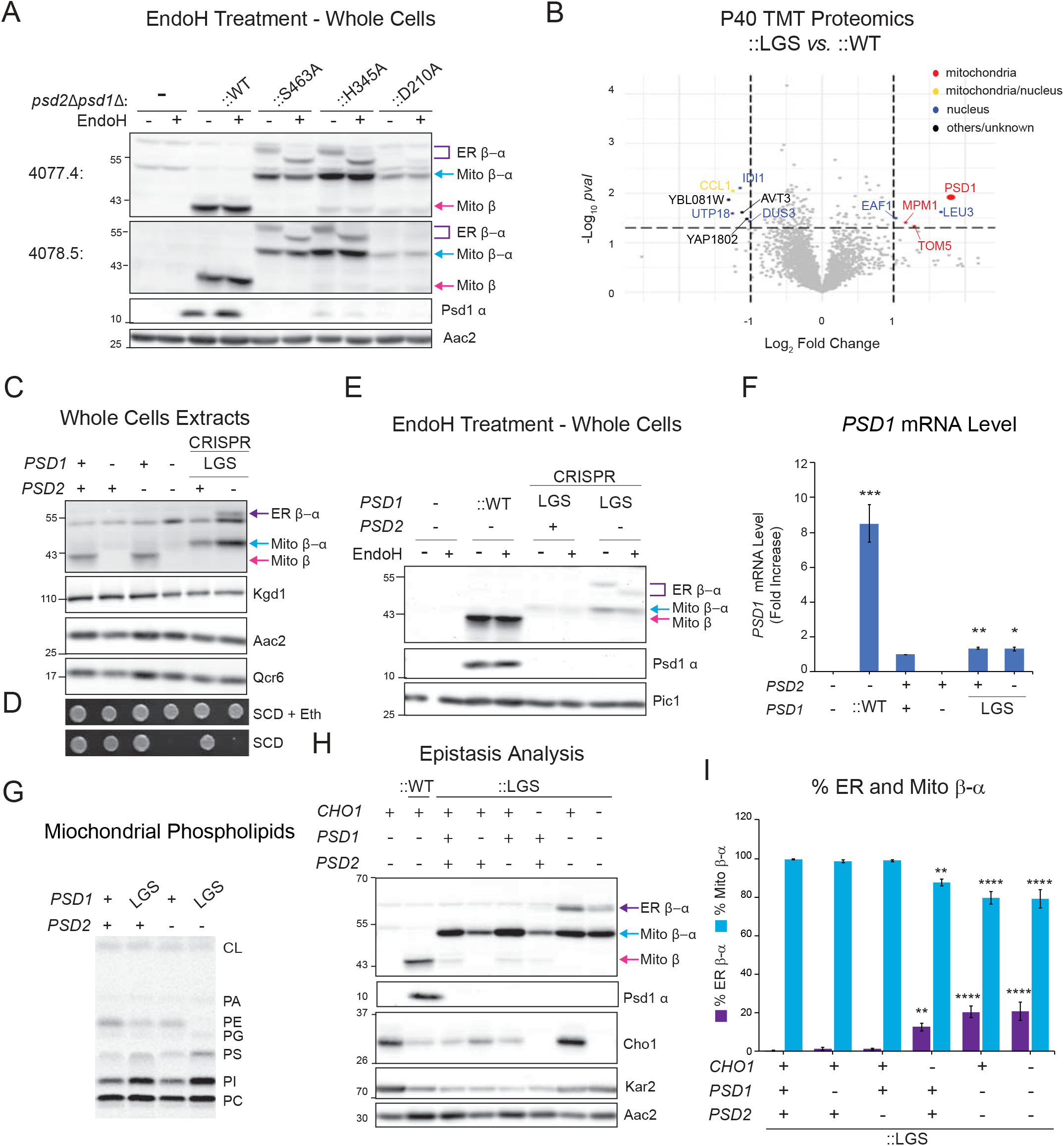
Glycosylation of endogenous and overexpressed mutant, nonfunctional Psd1 requires severe disruption of cellular PE metabolism. (A) Cell extracts from the indicated strains grown at 30°C in YPD were treated with EndoH as listed analyzed by immunoblot using the designated anti-Psd1 antisera; Aac2 served as loading control. ER β-α, glycosylated mutant Psd1; Mito β-α, mutant Psd1 not glycosylated (n=3). (B) TMT comparison of ER (P40) proteomes from *psd2*Δ*psd1*Δ∷LGS (n=3 preps) and *psd2*Δ*psd1*Δ∷WT yeast (n=4 preps). LGS; autocatalytic mutant Psd1. (C) Cell extracts from the indicated strains grown at 30°C in YPD were immunoblotted for the Psd1 β subunit; Kgd1, Aac2, and Qcr6 served as loading controls. LGS was knocked into endogenous *PSD1* of the listed strains via Hi-CRISPR (n=3). (D) Serial dilutions of same strains as in (C) were spotted onto SCD with or without 2 mM ethanolamine (+Eth) and incubated at 30°C for 3 days (n=3). (E) EndoH sensitivity assay of cell extracts from indicated strains grown in YPD at 30oC. Immunoblots were probed for Psd1 β (4077.4) and Psd1 α (FLAG); Pic1 served as loading control (n=3). (F) *PSD1* mRNA level in the indicated strains determined by two-step reverse transcription-quantitative PCR of *PSD1* and normalized to *ACT1* (means ± SEM, n=4). Statistical differences (1 symbol, P ≤ 0.05; 2 symbols P ≤ 0.01; 3 symbols P ≤ 0.001 compared WT (PSD2 +, PSD1 +) were calculated with unpaired Student’s t-test. (G) Mitochondrial phospholipids from the indicated strains were labeled overnight in YPD medium spiked with ^14^C-Acetate and separated by TLC (n=6). (H) Cell extracts derived from the indicated strains grown at 30°C in YPD were analyzed by immunoblot for Psd1 β (4077.4), Psd1 α (FLAG), Cho1 (PS synthase), and Kar2 (ER chaperone); Aac2 acted as loading control (n=5). (I) The relative abundance of ER β-α and Mito β-α in LGS mutant Psd1 expressing yeast of the indicated genotype was determined (means± SEM; n=5). Statistical differences (2 symbols P ≤ 0.01; 4 symbols P ≤ 0.0001) were determined by one-way analysis of variance (ANOVA) with Dunnett’s multiple comparison test.

Next, we performed a TMT proteomics comparison of the microsome-enriched P40 fractions from ∷WT versus those purified from *psd2*Δ*psd1*Δ∷Psd1-LGS/AAA (noted as ∷LGS; also has C-terminal 3XFLAG). Psd1, which was not elevated in P40 fractions from ∷WT versus those isolated from Psd-lacking yeast (Figure 1E), was detected at significantly increased levels in ∷LGS versus ∷WT microsome-enriched fractions (Figure 3B). Consistently, cell fractionation analysis of ∷LGS showed ER β-α in all fractions except the cytosol/S40 (Figure S3A). Notably, the unprocessed Psd1 Mito β-α band from ∷LGS had a similar fractionation profile to Mito β from ∷WT with each being highly enriched in the Qcr6-containing mitochondrial P13 fraction. As only two additional mitochondrial proteins (Mpm1 and Tom5) were significantly elevated in the ∷LGS microsomes (Figure 3B), we conclude that the accumulation of Psd1-LGS in the ER is specific to this nonfunctional form of Psd1.

Since overexpression of WT Psd1 induces the mito-CPR import stress response (Weidberg and Amon, 2018), we considered the possibility that the accumulation of glycosylated autocatalytic mutants of Psd1 was a by-product of their overexpression. Therefore, we used homology-integrated (HI) Clustered Regularly Interspaced Short Palindromic Repeats (CRISPR) to engineer two strains with the LGS/AAA mutation inserted into the *PSD1* gene. This mutation was edited in strains with and without *PSD2*. Consistent with *psd2*Δ*psd1*Δ strains transformed with autocatalytic Psd1 mutants (Figure 3A), two unprocessed forms of Psd1-LGS were detected in the absence of *PSD2*, Mito β-α and ER β-α, the latter of which was EndoH-sensitive (Figures 3C, E). Surprisingly, when *PSD2* was present, only one unprocessed form of Psd1-LGS was detected, Mito β-α, which was not sensitive to glycosidase (Figures 3C, E). Importantly, in the absence of *PSD2*, Psd1-LGS failed to support growth without ethanolamine supplementation (Figure 3D). We noticed that the steady state level of Psd1-LGS was increased in the absence versus the presence of *PSD2* (Figure 3C). Since *PSD1* mRNA levels were similar between these two strains (Figure 3F), the basis for this difference is post-transcriptional. As expected (Calzada et al., 2019), mitochondrial PE was reduced or essentially absent when Psd1-LGS was expressed in the presence or absence of *PSD2*, respectively (Figure 3G; Figure S3D).

Our failure to detect the ER β-α form of Psd1-LGS in the context of a functional Psd2, suggested that its accumulation may require severe perturbation of cellular PE metabolism. Consistent with this possibility, when Psd1-LGS was overexpressed in an otherwise WT yeast strain (*PSD1*+, *PSD2*+), only one unprocessed form of Psd1-LGS was detected which co-fractionated with the Qcr6- and endogenous Psd1-containing mitochondrial P13 fraction (Figure S3A). Interestingly, when expressed in WT yeast, Psd1-LGS was resistant to protease in intact mitochondria and only became accessible upon osmotic disruption of the OM or addition of detergent (Figure S3B). In contrast, when expressed in *psd2*Δ*psd1*Δ yeast, a noticeable amount of Psd1-LGS was resistant to protease upon OM-rupture (Figure S3B) and which was not related to any changes in its aggregation status (Figure S3C).

To directly test if the accumulation of autocatalytic mutant ER β-α depends on a severely-compromised phospholipid metabolism, we performed an epistasis analysis in which Psd1-LGS was integrated into strains lacking Psd1, Psd2, or Cho1, singly or in combination (Figure 3H). *CHO1* encodes the ER-resident PS synthase (Vance, 1990) that provides the substrate, PS, that is decarboxylated to PE by Psd1 and Psd2. Of consideration is that the *cho1*Δ*psd2*Δ*psd1*Δ∷LGS strain was generated through Hi-CRISPR based deletion of *CHO1* in the *psd2*Δ*psd1*Δ∷LGS parental strain; as such the abundance of Psd1-LGS forms in these strains are directly comparable. As observed for LGS knock-in strains (Figure 3C), glycosylated ER β-α was not detected in the absence of Psd1 (*psd1*Δ) or Psd2 (*psd2*Δ) alone, but was detected in their combined absence (Figures 3H, I). Further, glycosylated ER β-α accumulated in the absence of Cho1, and consistent with its role upstream of Psd1 and Psd2, the amount of ER β-α was not further increased upon its deletion in *psd1*Δ*psd2*Δ yeast (Figures 3H, I). These results indicate that the accumulation of the glycosylated, non-functional Psd1 precursor in the ER requires severe disturbance in cellular PE metabolism.

### Additional autocatalytic Psd1 mutants are also glycosylated

We screened a series of 33 alanine mutants of evolutionarily conserved residues by fluorescence immunoblotting with the goal of identifying additional mutations that either yield glycosylated forms of Psd1, regardless of their impact on autocatalysis, or that impair autocatalysis but are not partially glycosylated (Figure 4A, B). In this assay, Psd1 constructs that have successfully performed autocatalysis form mature Psd1 β (*red*) and Psd1 α (*green*) subunits that migrate separately, as for WT Psd1. In contrast, any mutation that impairs or ablates autocatalysis results in unprocessed forms of Psd1 that are co-detected by the Psd1 β and Psd1 α antisera (*yellow/orange*), as observed for autocatalytic null Psd1-LGS. For Psd1 mutants in which autocatalysis was not impeded, only a single Psd1 β band was detected that was not recognized by the Psd1 α antibody. Thus, no mutants that performed autocatalysis were glycosylated. We identified several additional mutations (R196A, R385A, 402-VGAT/AAAA-405, and 407-VGSI/AAAA-411) beyond the autocatalytic triad (D230, H345, and S463) that disrupted autocatalysis, each of which generated both unprocessed forms of Psd1, Mito β-α and ER β-α (Figures 2A, B). Intriguingly, three mutations (R189A, K230A, and H346A) that only partially impaired autocatalysis generated all three forms of Psd1, Mito β (*pink arrow*), Mito β-α (*blue arrow*) and ER β-α (*purple arrow*). It is worth mentioning that the co-detected (*yellow*) Psd1 bands migrating below Mito β are proteolytic fragments derived from unprocessed Mito β-α that are generated by mitochondrial proteases and accumulate when autocatalysis is impaired (Ogunbona et al., 2017). From this screen, we conclude that in the context of impaired cellular PE metabolism, any mutation that perturbs autocatalysis results in the accumulation of glycosylated ER β-α.

**Figure 4.**
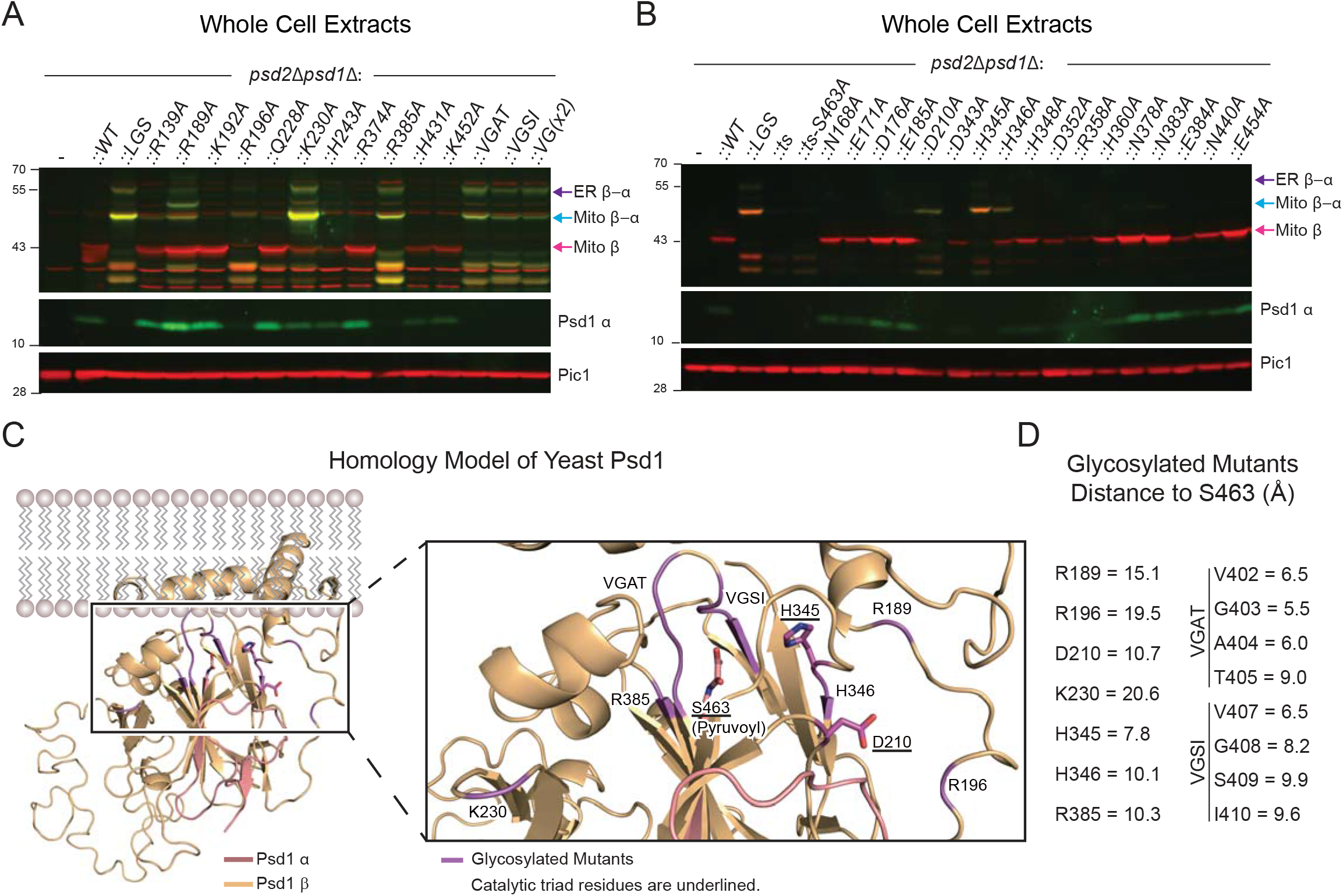
Mutations that indirectly impair autocatalysis are also glycosylated. (A, B) Cell extracts derived from the indicated strains grown at 30°C in YPD were analyzed by flourescent immunoblot for Psd1 β (red) and Psd1 α (green); Pic1 acted as loading control (n=3). (C) Homology model structure of Psd1 based on the E. coli PSD structure (PDB code: 6L06)—view from the side of membrane. The α subunit and β subunit are colored in light orange and salmon pink, respectively. The right panel is a magnified view around S463 (pyruvoyl group at N-terminus of α subunit). Residues whose mutation to alanine impairs autocatalysis are indicated and colored in magenta. Autocatalytic triad residues are underlined and shown in stick form. (D) The distances were measured between the carbonyl carbon of the pyruvoyl group (S463) and the alpha carbon of the residues whose mutation to alanine impairs autocatalysis.

To gain further insight, we built a homology model of yeast (*Saccharomyces cerevisiae*) Psd1 based on the *Escherichia coli* crystal structure (PDB code: 6L06) (Watanabe et al., 2020) using SWISS-MODEL (Figure 4C). Mutations that were not compatible with autocatalysis cluster in the immediate vicinity of S463 (cleavage occurs between G462 and S463). Similar to D210 (10.7 Å) and H345 (7.8 Å), H346 (10.1 Å), R385 (10.3 Å), VGAT (5.5-9.0 Å), and VGSI (6.5-9.9Å) are in close proximity to S463, the residue onto which the pyruvoyl prosthetic group is attached post-autocatalysis (Figure 4D). On the other hand, R189, R196, and K230 are greater than or equal to 15 Å away from S463 (Figure 4D). As R189 and R196 are close to the catalytic triad residues H345 and D210, respectively (Figure 4C), we speculate that their mutation to alanine may secondarily impact the positioning of these important residues that form the base and acid of the triad. K230, whose mutation also incompletely impairs autocatalysis (Figure 4A), is relatively distant from the catalytic site and thus may contribute to structural stabilization of an autocatalytic-competent form of the Psd1 precursor.

### Supplements of phospholipid synthesizing pathways decrease glycosylated mutant Psd1

Next, we asked if supplements that feed PE-biosynthetic pathways outside of the mitochondrion could prevent the accumulation of the glycosylated, non-functional mutant Psd1 protein in the ER. Indeed, supplementation of YPD with either ethanolamine, which promotes PE production via the ER-resident Kennedy pathway, or lyso-PE (LPE), which is converted to PE by Ale1, significantly diminished the relative amount of the glycosylated precursor protein (ER β-α; Figure 5A, B). LPE supplementation, which was shown to fully rescue mitochondrial defects when Psd1 is absent (Riekhof et al., 2007), decreased the detection of both Mito β-α and ER β-α precursor bands to a stronger degree than ethanolamine supplementation, which incompletely rescues the mitochondrial defects when Psd1 is missing (Calzada et al., 2019).

**Figure 5.**
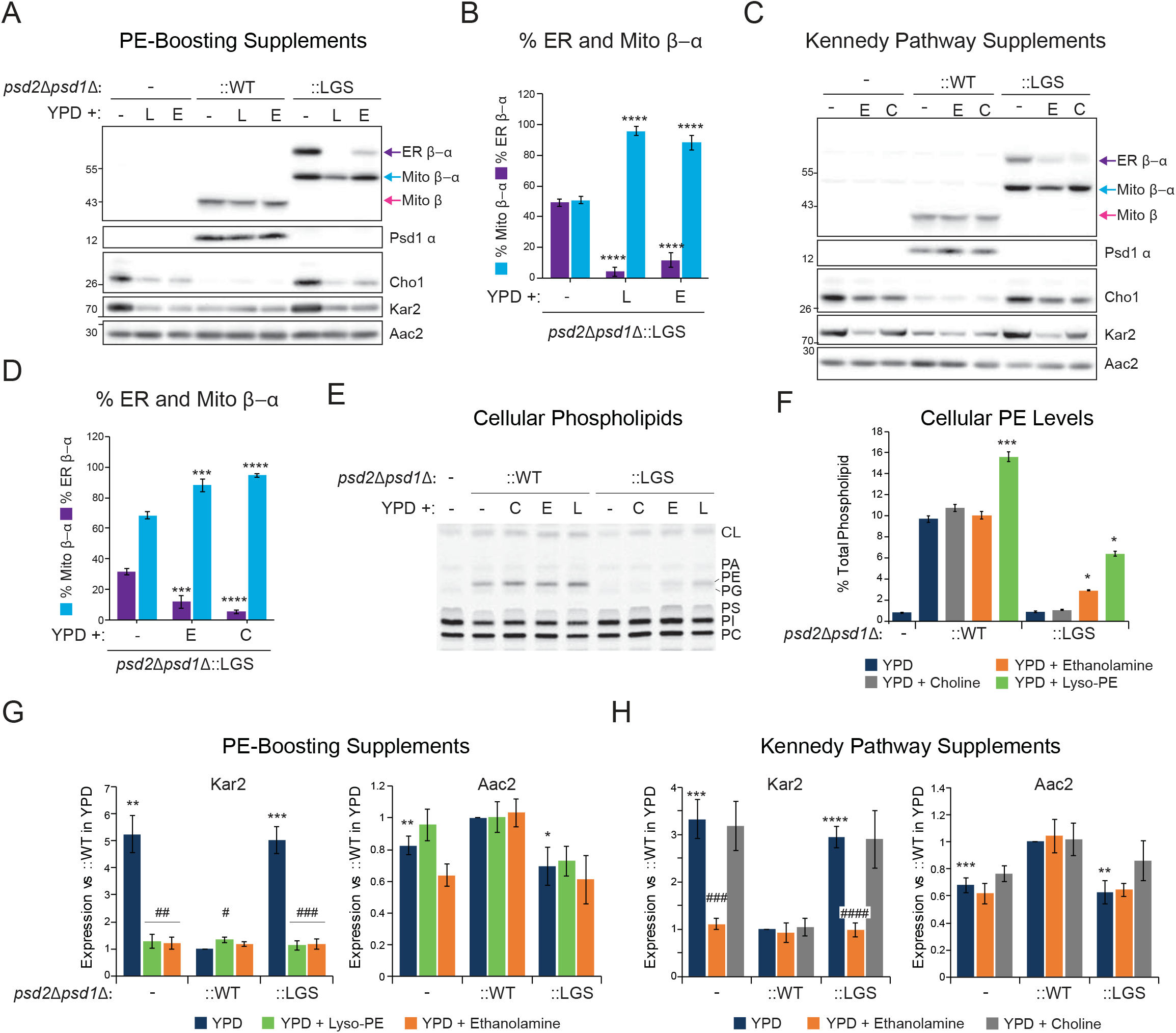
Supplements for distinct ER-resident phospholipid biosynthetic pathways decrease accumulation of glycosylated nonfunctional Psd1. (A) Pre-cultures of the indicated strains were inoculated in YPD medium supplemented with 1% (v/v) Tergitol (-), 0.5mM lyso-phosphatidylethanolamine in 1% (v/v) Tergitol (L), or 2mM ethanolamine (E). Following overnight growth at 30oC, cell extracts were harvested and immunoblotted for Psd1 β, Psd1 α, Cho1, and Kar2; Aac2 acted as loading control (n=5). (B) The relative abundance of LGS mutant Psd1 ER β-α and Mito β-α was determined from yeast analyzed in (A) (mean ± SEM, n=5). (C) Pre-cultures of the indicated strains were inoculated in YPD medium alone or supplemented with 2mM ethanolamine (E) or 2mM choline (C). Following overnight growth at 30oC, cell extracts were harvested and immunoblotted as listed. (D) The relative abundance of LGS mutant Psd1 ER β-α and Mito β-α was determined from yeast analyzed in (C) (mean ± SEM, n=5). For B and D statistical differences (3 symbols P ≤ 0.001; 4 symbols P ≤ 0.0001) compared to *psd2*Δ*psd1*Δ were calculated by one-way ANOVA with Dunnett’s multiple comparison test. (E) Cellular phospholipids from the indicated strains grown in YPD alone or supplemented with choline, ethanolamine, or lyso-phosphatidylethanolamine were labeled overnight with ^14^C-Acetate and separated by TLC (n=6). (F) Cellular PE abundance was determined for each strain in each growth condition (mean ± SEM for n=6). Statistical differences (1 symbol P ≤ 0.05; 3 symbols P ≤ 0.001) relative to growth in YPD alone was determined by one-way ANOVA with Holm-Sidak pairwise comparison (for ∷WT samples) or one-way ANOVA by Ranks (∷LGS samples). (G) Steady-state abundance of Kar2 and Aac2 in indicated strains grown in absence or presence of lyso-PE or ethanolamine relative to *psd2*Δ*psd1*Δ∷WT yeast grown in YPD alone (mean ± SEM for n=4). (H) Steady-state abundance of Kar2 and Aac2 in indicated strains grown in absence or presence of ethanolamine or choline relative to *psd2*Δ*psd1*Δ∷WT yeast grown in YPD alone (mean ± SEM for n=5). For G and H, statistical differences (1 symbol P ≤ 0.05; 2 symbols P ≤ 0.01; 3 symbols P ≤ 0.001; 4 symbols P ≤ 0.0001) compared to WT in YPD alone (*) or YPD alone for a given strain (#) were calculated with unpaired Student’s t-test.

To determine if the ability of lyso-PE and ethanolamine to decrease the relative amount of ER β-α was specific to ER-resident PE biosynthetic pathways, we used choline, which promotes PC biosynthesis in the ER (Walkey et al., 1998). Surprisingly, the addition of choline also decreased the proportion of nonfunctional ER β-α (Figure 5C, D). Lipid analyses confirmed that cellular PE levels were significantly increased when cultures were supplemented with LPE and ethanolamine but not choline (Figure 5E, F). Kar2, a marker of ER stress (Hsu et al., 2012; Lajoie et al., 2012), was increased in *psd2*Δ*psd1*Δ, as seen previously (Calzada et al., 2019), and ∷LGS relative to ∷WT (Figure 5G, H). Interestingly, the addition of LPE or ethanolamine, but not choline, to YPD medium drastically reduced steady state Kar2 levels in both *psd2*Δ*psd1*Δ and ∷LGS yeast. In combination, these results indicate that supplements that feed distinct ER-resident lipid biosynthetic pathways reduce the accumulation of glycosylated non-functional Psd1-LGS by a mechanism(s) that is not strictly related to their capacity to promote PE production.

### Glycosylated Psd1 mutant is short-lived and ubiquitinated

We reasoned that the ability of the ER lipid-boosting supplements to reduce the steady state amount of ER β-α could reflect their ability to decrease its generation or alternatively, increase its removal. As such, we performed cycloheximide-based *in vivo* degradation assays to determine the relative stabilities of both unprocessed forms of Psd1-LGS in comparison to the separated α and β subunits of WT Psd1, and whether or not choline exerted any affect (Figure 6A). In the absence of choline, the WT Psd1 α and β subunits were stable for the 2hr incubation following inhibition of cytosolic translation with cycloheximide (Figure 6A, B), as previously observed (Ogunbona et al., 2017). In contrast, the steady state amounts of ER β-α and Mito β-α from Psd1-LGS each decreased significantly over time, and strikingly, ER β-α decreased much faster than Mito β-α (Figure 6A, C). Moreover, the relative stability or instability of WT Psd1 and Psd1-LGS was not impacted by choline supplementation. Given that choline did not significantly alter *PSD1-LGS* transcript levels (Figure S4A), this suggests that choline supplementation decreases ER β-α abundance by a post-transcriptional mechanism that is not related to ER stress (Figure 5H) and which does not reflect its ability to stimulate ER β-α degradation (Figure 6C).

**Figure 6.**
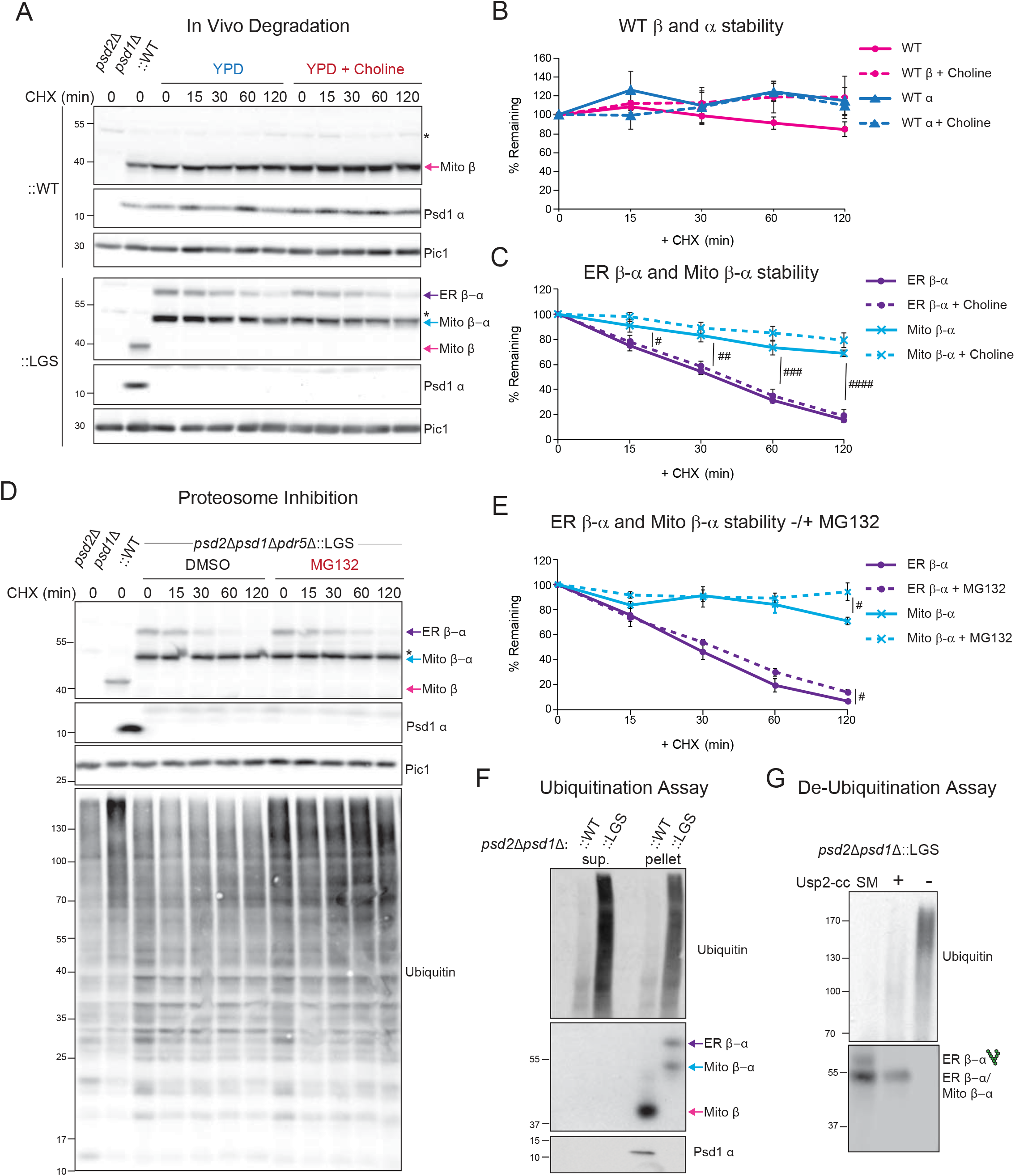
Glycosylated non-functional mutant Psd1 is short-lived and ubiquitinated. (A) *In vivo* degradation assay. Cell extracts from designated strains were isolated at the indicated times following growth in YPD containing cycloheximide (CHX) only or CHX and choline. Samples were resolved by SDS-PAGE and immunoblotted as indicated (n=5). *, nonspecific bands. (B) The percentages of WT α and β subunits remaining at each time point were quantified (mean ± SEM for n = 5). (C) The percentages of nonfunctional glycosylated ER β-α and mitochondrial β-α remaining at each time point were quantified (mean ± SEM for n = 5). Statistical differences (1 symbol P ≤ 0.05; 2 symbols P ≤ 0.01; 3 symbols P ≤ 0.001; 4 symbols P ≤ 0.0001) between ER β-α and mito β-α was determined at each timepoint by unpaired student t-test. (D) An overnight YPD culture of the *psd2*Δ*psd1*Δ*pdr5*Δ∷LGS strain was resuspended in SCD with 2mM choline and further spiked with vehicle (DMSO) or the proteosomal inhibitor MG132. After a 1hr incubation at 30°C, CHX was added and cell extracts harvested following growth at 30oC for the indicated times. Samples were resolved by SDS-PAGE and immunoblotted as indicated (n=5). (E) The percentages of nonfunctional glycosylated ER β-α and mitochondrial β-α remaining at each time point were quantified (mean ± SEM for n = 5). Statistical differences (1 symbol P ≤ 0.05) between vehicle and MG132 treated samples for ER β-α and mito β-α was determined at each timepoint by unpaired student t-test. (F) *In vivo* ubiquitination assay. Crude lysate was prepared from the indicated strains and lysates were ultracentrifuged into soluble and membrane fractions, the latter of which was solubilized with digitonin. FLAG tagged WT and mutant Psd1 was immunoprecipitated from the soluble and digitonin-extracted membrane fractions and Mito β, Mito α, ER β-α and Mito β-α detected by immunoblot using 4077.4 antisera; ubiquitin antibody detected ubiquitination (n=3). (G) *In vivo* re-translocation assay. Ubiquitin removal with Usp2Core, quenched with SUME, immunoprecipitated with anti-FLAG resin, and immunoblotted for Psd1 β and ubiquitin (n=3).

Given its atypical localization and comparatively short half-life, we postulated that the cytosolic proteasomal system may be responsible for the rapid degradation of ER β-α. As such, we cultured ∷LGS cells lacking the multi-drug exporter Pdr5 (*psd2*Δ*psd1*Δ*pdr5*Δ∷LGS) in SCD medium supplemented with 2 mM choline (included to support growth of *psd2*Δ*psd1*Δ which are auxotrophs for ethanolamine or choline (Atkinson et al., 1980; Birner et al., 2001; Burgermeister et al., 2004)) in the absence (DMSO) or presence of the proteasomal inhibitor MG132. While the effect was modest, MG132 slightly stabilized both ER β-α and Mito β-α following proteasomal inhibition for 2hrs (Figure 6D, E). As expected, ubiquitinated proteins accumulated when ∷LGS yeast were treated with MG132 but not DMSO (Figure 6D). The fact that Mito β-α was stabilized by MG132 to a similar extent as ER β-α suggests a role for the cytosolic proteasomal system in the turnover of both unprocessed forms of Psd1-LGS. The transcript levels for numerous plasma membrane ATP binding cassette transporters were significantly increased in ∷LGS versus ∷WT (Figure S4B). As such, we speculate that the relatively weak ability of MG132 to stabilize Psd1-LGS could reflect the increased expression of multi-drug pumps in this yeast strain.

Proteasomal degradation of a subset of substrates requires removal from their organellar homes― in a process known as retro-translocation― for degradation by the cytosolic proteasome (Neal et al., 2018). As such, we examined the ubiquitination status of Psd1-LGS compared to WT Psd1 and analyzed whether membrane-residing ubiquitinated Psd1-LGS retro-translocated to the cytosol for proteasomal degradation (Figure 6F). Cell extracts were ultracentrifuged to separate the soluble cytosol from membrane-containing organelles, the latter of which was then solubilized with digitonin, a mild detergent that preserves the non-covalent association of Psd1 α and β subunits (Ogunbona et al., 2017). Next, Psd1 was immunoprecipitated from each fraction using anti-FLAG conjugated beads and bound material analyzed by immunoblot. While both subunits of WT Psd1 and both unprocessed forms (ER β-α and Mito β-α) of Psd1-LGS were immunoprecipitated from digitonin-extracted membrane pellets, only the anti-FLAG bound material from Psd1-LGS was robustly ubiquitinated (Figure 6F). The associated ubiquitin signal was even stronger following FLAG immunoprecipitation of Psd1-LGS from the cytosolic supernatant fractions, even though neither ER β-α or Mito β-α was detected by the Psd1 antibody (Figure 6F). To confirm that the ubiquitin signal resulted from Psd1-LGS, the supernatant fraction was incubated with human recombinant Usp2Core, a broadly active ubiquitin protease, prior to FLAG immunoprecipitation (Figure 6G). Following Ups2 pretreatment, the ubiquitin signal was decreased and a single unprocessed form of Psd1-LGS observed. These data indicate that under standard growth conditions, there is a measurable amount of retro-translocated non-functional mutant Psd1-LGS in the cytosol that has been deglycosylated and is en route to the proteasome for degradation.

### Supplements that diminish glycosylated mutant Psd1 act by distinct mechanisms

When grown in dextrose, the expression of many mitochondrial genes is repressed, including those involved in oxidative phosphorylation, and the vast majority of cellular energy is generated by glycolysis. One consequence of a less active electron transport chain is a proportionally weaker membrane potential across the IM which serves as a critical driving force for import and assembly of mitochondrial precursors destined for the IM or matrix. Therefore, we asked whether the relative proportion of the Mito β-α and ER β-α unprocessed forms of Psd1-LGS are influenced by metabolic growth condition (Figure S4C). Indeed, the relative abundance of ER β-α was significantly decreased, and Mito β-α proportionately increased, following growth in YPEG, which requires mitochondrial function (Figures S4 A-C). However, in rich lactate, which contains a small amount of dextrose, ER β-α and Mito β-α were only moderately altered.

Yeast lacking Psd1 have a respiratory growth defect that is at least in part attributed to a significant reduction in respiratory complex IV activity and a modest decline in complex III function (Calzada et al., 2019). In the combined absence of Psd1 and Psd2, there is an additive effect such that the *psd1*Δ respiratory growth phenotype is exacerbated and the activities of respiratory complexes III and IV are almost completely ablated (Calzada et al., 2019). Therefore, we reasoned that the inability to detect ER β-α when Psd2 is expressed (Figure 3H) could reflect an increased IM membrane potential secondary to this strain’s higher respiratory capacity. To test this idea, we utilized the protonophore carbonyl cyanide *m*-chlorophenyl hydrazone (CCCP), a pharmacological uncoupler that collapses the proton gradient across the IM which in turn blocks protein import into and across this membrane. In the presence of Psd2 but absence of CCCP, only one unprocessed form of Psd1-LGS, Mito β-α, was detected (Figure 7A). However, in presence of Psd2 and CCCP, a small amount of ER β-α accumulated (Figure 7A), indicating that severe import stress can stimulate the accumulation of glycosylated Psd1-LGS. Notably, only a single, fully processed Mito β was detected for WT Psd1 in either the absence or presence of IM uncoupling.

**Figure 7.**
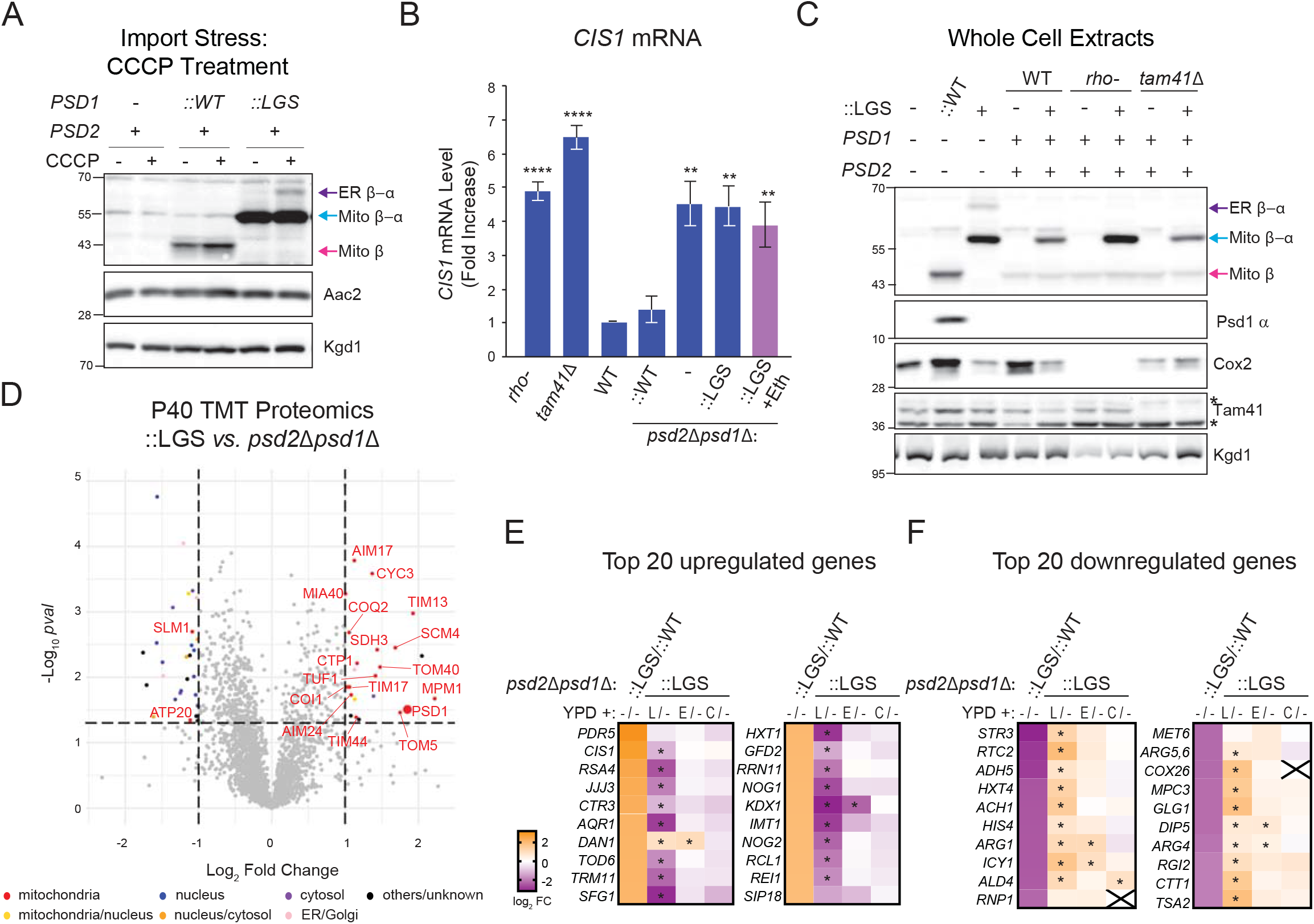
Supplements that diminish amount of glycosylated mutant Psd1 act by distinct mechanisms. (A) The indicated strains were cultured overnight at 30 °C in YPD without or with 10 μM CCCP. Cell extracts were harvested and immunoblotted as designated (n=3). (B) *CIS1* mRNA levels in the indicated strains grown in YPD at 30°C were determined by two-step reverse transcription-quantitative PCR and normalized to *ACT1* (mean ± SEM, n=5). Statistical differences (2 symbols P ≤ 0.01; 4 symbols P ≤ 0.0001) versus WT were calculated with unpaired Student’s t-test. (C) Cell extracts from the indicated strains, untransformed or transformed with the autocatalytic LGS Psd1 mutant, grown overnight in YPD were immunoblotted for Psd1 β (4077.4), Psd1 α (FLAG), Cox2 (encoded by mtDNA), and Tam41; Kgd1 acted as loading control (n=3). *, nonspecific bands. (D) TMT comparison of ER (P40) proteomes from *psd2*Δ*psd1*Δ∷LGS and *psd2*Δ*psd1*Δ yeast (n=3 preps each). LGS; autocatalytic mutant Psd1. (E and F) Heatmaps of gene expression analysis from RNAseq with the indicated strains grown in YPD medium alone, or YPD supplemented with lyso-PE (L), ethanolamine (E), or choline (C). padj values that were significantly different from YPD medium alone (P ≤ 0.05) are designated (*). (E) The top 20 upregulated genes in ∷LGS vs ∷WT and how each supplement does or does not affect their expression relative to ∷LGS grown in YPD alone. (F) The top 20 downregulated genes in ∷LGS vs ∷WT and how each supplement does or does not affect their expression relative to ∷LGS grown in YPD alone.

It was previously demonstrated that mitochondrial import stress activates the mito-CPR pathway whose hallmark feature is an increase in *CIS1* mRNA levels (Weidberg and Amon, 2018). Thus, we asked whether the glycosylated Psd1-LGS mutant induces *CIS1* transcription. As previously reported (Weidberg and Amon, 2018), *CIS1* transcript levels, which are normally low, were elevated in *rho-* and *tam41*Δ yeast (Figure 7B), strains with mitochondrial import defects stemming from the loss of mitochondrial DNA or defective cardiolipin biosynthesis, respectively. Compared to WT yeast, *CIS1* mRNA levels were also significantly increased in *psd2*Δ*psd1*Δ and ∷LGS strains but not in the *psd2*Δ*psd1*Δ strain rescued with WT Psd1 (∷WT). These results indicate that mito-CPR is activated, likely due to impaired mitochondrial protein import, when cellular PE metabolism is severely compromised regardless of whether or not a nonfunctional form of Psd1 is expressed. Surprisingly, *CIS1* transcript levels were not reduced when ∷LGS yeast were supplemented with ethanolamine (Figure 7B, *pink bar*). This implies that ethanolamine reduces the relative abundance of ER β-α (Figure 5A) in a mito-CPR independent manner. Consistent with this interpretation, when Psd1-LGS was integrated into *rho-* and *tam41*Δ yeast, only a single unprocessed form, Mito β-α, was detected (Figure 7C). From these results, we conclude that mito-CPR is not itself sufficient to drive the accumulation of glycosylated nonfunctional Psd1 mutants.

As mito-CPR was similarly activated in the *psd2*Δ*psd1*Δ and ∷LGS strains and unaffected by ethanolamine, we next asked whether the expression of the Psd1-LGS mutant imposes any overt changes beyond that typically observed in the already severely PE-deficient *psd2*Δ*psd1*Δ strain. To this end, we compared the microsome-enriched P40 proteomes of ∷LGS versus *psd2*Δ*psd1*Δ (Figure 7D). This comparison showed that numerous mitochondrial proteins in addition to Psd1 were significantly increased in the ∷LGS versus *psd2*Δ*psd1*Δ P40 fractions, indicating that additional mitochondrial proteins are mislocalized. Given that non-Psd1 mitochondrial proteins were not significantly enriched in microsomes from either *psd2*Δ*psd1*Δ (Figure 1E) or ∷LGS (Figure 3B) relative to ∷WT, these results raise the possibility that the combination of compromised cellular PE metabolism and expression of Psd1-LGS, which is both difficult to import and non-functional, impinge upon mitochondrial biogenesis in distinct but partially overlapping manners.

To gain additional insight into the molecular underpinnings for how the ER lipid-boosting supplements reduce the steady state amount of ER β-α, we determined their impact on the transcriptomes of ∷WT and ∷LGS yeast (Figure 7E, F). When grown in dextrose alone, *PDR5*, which encodes the major drug pump, and the mito-CPR marker *CIS1* were the two most upregulated genes in ∷LGS versus ∷WT (Figure 7E). Additional genes in the Mitoprotein-Induced Stress Response (Boos et al., 2019; Boos et al., 2020), including *RPN4* and *PDR3* that function upstream of *CIS1*, were also significantly upregulated in ∷LGS versus ∷WT (Figure S4F). Many of the other top upregulated genes in dextrose-grown ∷LGS have roles related to cytosolic translation (e.g. *RSA4*, *TOD6*, *TRM11*, *RRN11*, *NOG1*, *NOG2*, *IMT1*, *RCL1*, and *REI1*). In contrast, many of the top downregulated genes have diverse functions in metabolism that include amino acid biosynthesis (*STR3*, *HIS4*, *ARG1*, *ARG4*, *ARG5,6*, and *MET6*), other biosynthetic processes (*ADH5* and *GLG1*), vacuolar amino acid transport (*RTC2* and *DIP5*), reactive oxygen detoxification (*TSA2* and *CTT1*), and mitochondrial fuel utilization *(MPC3*) and efficiency (*COX26*) (Figure 7F). Many, including *CIS1*, but not all, of these changes were corrected when ∷LGS yeast were grown with LPE (Figures 7E, F). In contrast, ethanolamine, which like LPE increased cellular PE levels (Figure 5F), albeit to a lesser extent, and reduced the amount of Kar2 (Figure 5G, H), was comparatively ineffective at normalizing the altered transcripts in ∷LGS. With one exception (*ALD4*), choline did not impact the altered transcriptional profile of ∷LGS yeast grown in dextrose. Overall, our combined results provide strong evidence supporting the conclusion that LPE, ethanolamine, and choline reduce the relative abundance of ER β-α by distinct mechanisms.

## DISCUSSION

Our data indicate that the vast majority, if not all, of functional Psd1 localizes to the mitochondrion. This conclusion is based on results from multiple yeast strain backgrounds, overexpression studies in yeast and mammalian cells, and a powerful combination of chimeric constructs and biochemical and immunofluorescence approaches. Further, it is consistent with the failure to detect significant Psd activity in microsomes when the non-mitochondrial Psd2 is missing (Ogunbona et al., 2017; Trotter and Voelker, 1995) and a recent high-confidence proteomics study, which failed to annotate Psd1 as being localized to organelles other than mitochondria (Morgenstern et al., 2017). Based on this overwhelming confluence, we conclude that a functionally relevant population of Psd1 does not localize to the ER, as recently suggested (Friedman et al., 2018). As such, the myriad of studies in which the strict mitochondrial localization of Psd1 served as a foundational premise (Baker et al., 2016; Birner et al., 2001; Bykov et al., 2020; Cui et al., 1993; Daum et al., 1986; Heden et al., 2019; Kornmann et al., 2009; Lahiri et al., 2014; Li and Dowhan, 1988; Steenbergen et al., 2005; Tamura et al., 2012a; Tasseva et al., 2013) do not need to be re-evaluated in consideration of a dual localization for this mitochondrial enzyme, as contended (Friedman et al., 2018).

While the final mature and functional enzyme localizes to mitochondria, there is clearly an interesting and perhaps unexpected relationship between Psd1 and the ER. First, proximity-specific ribosome profiling indicates that Psd1 translation occurs in the vicinity of both mitochondria and the ER, similar to Osm1, a protein that is dually localized to these two compartments (Williams et al., 2014). In fact, this observation motivated the study by (Friedman et al., 2018). Second, when expressed in yeast with severely compromised cellular PE metabolism, autocatalytic Psd1 mutants do engage the ER where they are glycosylated and quickly degraded. Together, these two observations suggest that some fraction of Psd1 biogenesis normally occurs in the context of the ER and when problems with its ability to be properly imported and fully matured arise, it can again be directed to the ER which now exerts a quality control functionality. Classically, mitochondrial precursors are viewed as being translated in the cytosol and, with the help of cytosolic chaperones, directly targeted to the mitochondrial translocase of the OM (TOM), the common entry gate for most nuclear-encoded mitochondrial proteins (Hansen and Herrmann, 2019). Recently, an ER-assisted pathway for a subset of mitochondrial proteins termed ER-SURF (ER-surface-mediated targeting) was discovered in yeast that functions in parallel to the classic direct import pathway (Hansen et al., 2018). Whether or not Psd1 is an ER-SURF substrate and if so, whether or not the components of this pathway, most notably Djp1, can discriminate between functional and non-functional Psd1 precursors are interesting questions for future studies. Further, it must be acknowledged that given this intimate relationship with the ER, there is the opportunity that there may be biological contexts and situations in which it is harnessed such that functional Psd1 is targeted to this organelle.

Although the effect was modest, proteasomal inhibition stabilized both unprocessed forms of Psd1-LGS to the same extent (Figure 6E). When considered in the context of the striking ubiquitination of Psd1-LGS in both membrane-associated and soluble cytosolic fractions, this strongly implicates the ubiquitin proteasome system in mediating the removal of both ER β-α and Mito β-α, although based on the relative kinetics of their degradation (Figure 6C), ER β-α is the easier substrate to resolve. The ubiquitin-proteasome system has a documented role in policing mitochondrial precursor biogenesis and ensuring that non-functional and/or mis-localized proteins are degraded. Recently, the mitochondrial protein translocation-associated degradation (mito-TAD) pathway was identified that monitors the routine import of precursors through TOM (Martensson et al., 2019). When precursor translocation is stalled, Ubx2, which also participates in the ER-associated degradation (ERAD) of misfolded proteins (Neuber et al., 2005), recruits Cdc48 together with its cofactors Npl4 and Ufd1 to deliver unwanted proteins to the proteasome for degradation (Martensson et al., 2019). When mitochondrial biogenesis is challenged, which according to our data occurs when cellular PE metabolism is knee-capped (Figure 7B), mito-CPR is activated to enhance the normal maintenance provided by mito-TAD (Weidberg and Amon, 2018). Mito-CPR induces the expression of Cis1 through Pdr3 to recruit the highly-conserved OM-anchored and cytosol-facing, AAA-ATPase Msp1 which functions as an extractase that removes clogged protein precursors from the TOM complex and targets them for proteasomal degradation directly or after first sending them to the ER (Matsumoto et al., 2019). It is tempting to speculate that when expressed in a PE-limited setting, Psd1-LGS stimulates Msp1 activity, which then contributes to the increased abundance of multiple mitochondrial proteins in microsomes from ∷LGS versus *psd2*Δ*psd1*Δ (Figure 7D). In this context, the fact that only one form of unprocessed Psd1-LGS, Mito β-α, was readily detected in the cardiolipin-deficient *tam41*Δ is particularly interesting (Figure 7C). While cardiolipin and cellular PE deficiency each causes import stress that activates mito-CPR, there are at least nuanced differences in how they are registered by the cell and in the responses that they generate. Ongoing experiments are being performed to dissect the contribution of these assorted quality control pathways and factors.

The ability of LPE and ethanolamine to diminish the accumulation of ER β-α in *psd2*Δ*psd1*Δ was initially not surprising as each compound feeds a different ER-resident PE biosynthetic pathway. Based on the literature and our prior work, the fact that LPE provided a bigger reduction in ER β-α than ethanolamine was taken to reflect its capacity to completely (Riekhof et al., 2007), instead of only partially as in the case of ethanolamine (Calzada et al., 2019), rescue the mitochondrial defects that occur in the absence of Psd1. The initial surprise was that choline, which we used as a negative control to demonstrate the specificity of the rescue to PE production, actually worked better in this regard than ethanolamine (Figure 5D). This was the first indication that the stress response activated by Psd1-LGS as expressed in a PE-deficient setting involved processes beyond PE and/or that can be reversed/overcome by PE-independent means. The sum of our results using these three supplements provide strong evidence that each reduces the amount of ER β-α through distinct mechanisms. The most likely mode of action for LPE is through its ability to significantly increase PE levels. If correct, then many of the transcriptional changes that occur in ∷LGS, especially those that are reversed by LPE, are the direct consequence of PE-depleted membranes. Ethanolamine, which rescued PE to ~50% of LPE levels, did decrease Kar2-associated ER stress to a similar degree as LPE. Thus, it is possible that the ability of ethanolamine and LPE to decrease ER β-α reflects their impact on ER stress. According to this possibility, their different capacities to revert the transcriptional changes in ∷LGS could indicate that a threshold level of PE was not achieved by ethanolamine but was with LPE. Recently, it was demonstrated that ethanolamine rescues mitochondrial respiration of cardiolipin mutant yeast by a PE-independent mechanism (Basu Ball et al., 2018). Thus, it is certainly possible that the underlying mechanism for the reduction provided by ethanolamine is only partially due to its ability to augment PE. The mechanism by which choline decreases ER β-α is the most mysterious and perhaps most interesting. Choline does not decrease Kar2-associated ER stress and in fact, may be expected to increase it (Thibault et al., 2012). Moreover, choline does not increase the rate of ER β-α disappearance (Figure 6C) nor reduce the elevated *CIS1* levels in ∷LGS. Moving forward, it will be exciting to mechanistically distinguish how these supplements reduce ER β-α accumulation in the context of cellular PE deficiency and perhaps more broadly in additional models of mitochondrial import stress.

## LIMITATIONS OF STUDY

The failure to detect a glycosylated form of WT Psd1 does not formally demonstrate that it does not exist either normally or in specific situations. While this is a noted limitation of this study, our results do forcefully demonstrate that a significant population of functional Psd1 is not likely to reside in the ER under the conditions employed here. Also, our work with autocatalytic Psd1 mutants was limited to yeast models. It will be important to ascertain whether some of the principles established in yeast are conserved in other eukaryotic systems. Finally, the ability of the ER-assisted adaptive response that is activated by dysfunctional cellular PE metabolism to detect other mitochondrial precursors was not formally tested beyond the proteomics comparison of ∷LGS and *psd2*Δ*psd1*Δ microsomal fractions (Figure 7D).

## RESOURCE AVAILABILITY

### Lead Contact

Requests for resources and reagents should be directed to and will be fulfilled by the Lead Contact, Steven Claypool.

### Materials Availability

All unique reagents and materials generated in this study are available from the Lead Contact without restriction, except for the possible completion of a Materials Transfer Agreement.

### Data and Code Availability

The RNASeq datasets are available at the Gene Expression Omnibus under accession number GSE162987.

The TMT proteomics dataset is available at the MassIVE repository massive.ucsd.edu and can be accessed using the link (ftp://MSV000086558@massive.ucsd.edu).

Original/source data for all figures presented in the paper are available from the Lead Contact upon request.

## METHODS

### Molecular Biology

All of the Psd1 constructs used in this study were subcloned into the pRS305 plasmid. The following constructs have been previously described: WT Psd1 containing a COOH-terminal 3XFLAG tag, the autocatalytic LGS/AAA mutant, and ER-Psd1 (Onguka et al., 2015); the Psd1 point mutants analyzed in Figures 3A and 4B (Ogunbona et al., 2017; Zhao et al., 2019); and human PISD with 3XFLAG tag (Zhao et al., 2019). Additional point mutations analyzed in Fig. 4A (R139A, R189A, K192A, R196A, Q228A, K230A, H243A, H374A, R385A, H431A, K452A, V402GAT405, V407GSI410, and VG(x2)) were generated by overlap extension using pRS305Psd3XFLAG as template. To direct Psd1 to the IM using information from other IM proteins with a similar topology, the first 101 amino acids of Psd1, which includes its mitochondrial targeting sequence and single transmembrane domain, were replaced by the equivalent regions of the following single pass IM proteins: Tim50 (1-132), Mic60 (1-57), Yme1 (1-251), Tim54 (1-54), and Ccp1 (1-36). All constructs were verified by DNA sequencing.

### Yeast strain generation and growth conditions

Yeast strains used in this study are listed in Table 1 and were derived from GA74-1A, BY-4741, W303 or D273-10B. To integrate assorted pRS305-based constructs into the yeast genome, AflII-linearized plasmids were transformed into GA74-1A yeast strains of the indicated genotype (in general, *psd1*Δ*psd2*Δ; however, the autocatalytic mutant LGS/AAA Psd1 mutant was additionally transformed into WT yeast containing or lacking mtDNA (*rho-*), and *psd1*Δ, *psd2*Δ, *cho1*Δ, and *tam41*Δ single deletion strains), stable integrants selected on synthetic dropout medium lacking Leu (0.17% yeast nitrogen base, 0.5% ammonium sulfate, 0.2% dropout mixture synthetic-leu, 2% dextrose), and expression of each Psd1 construct verified by immunoblot.

**TABLE 1.**
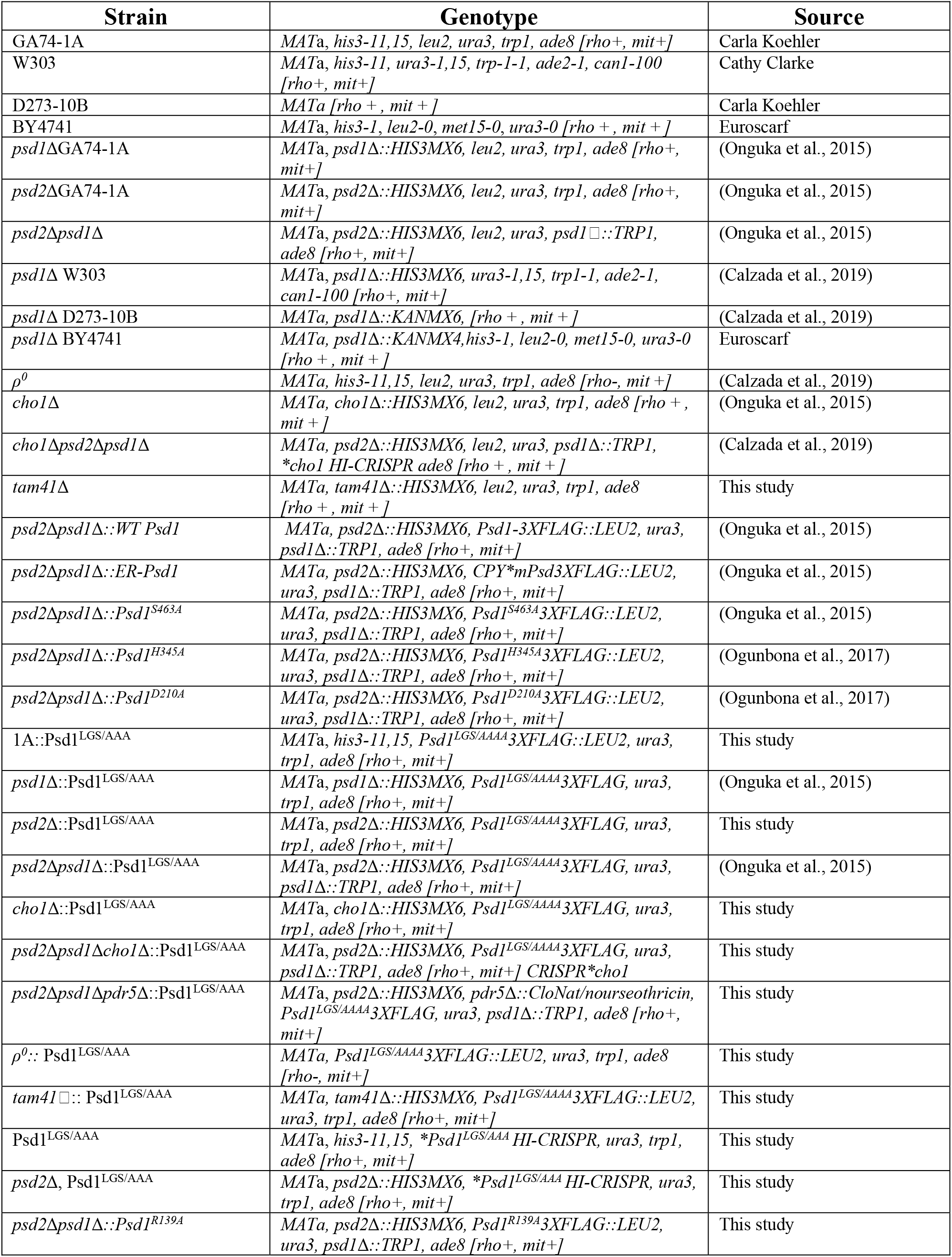
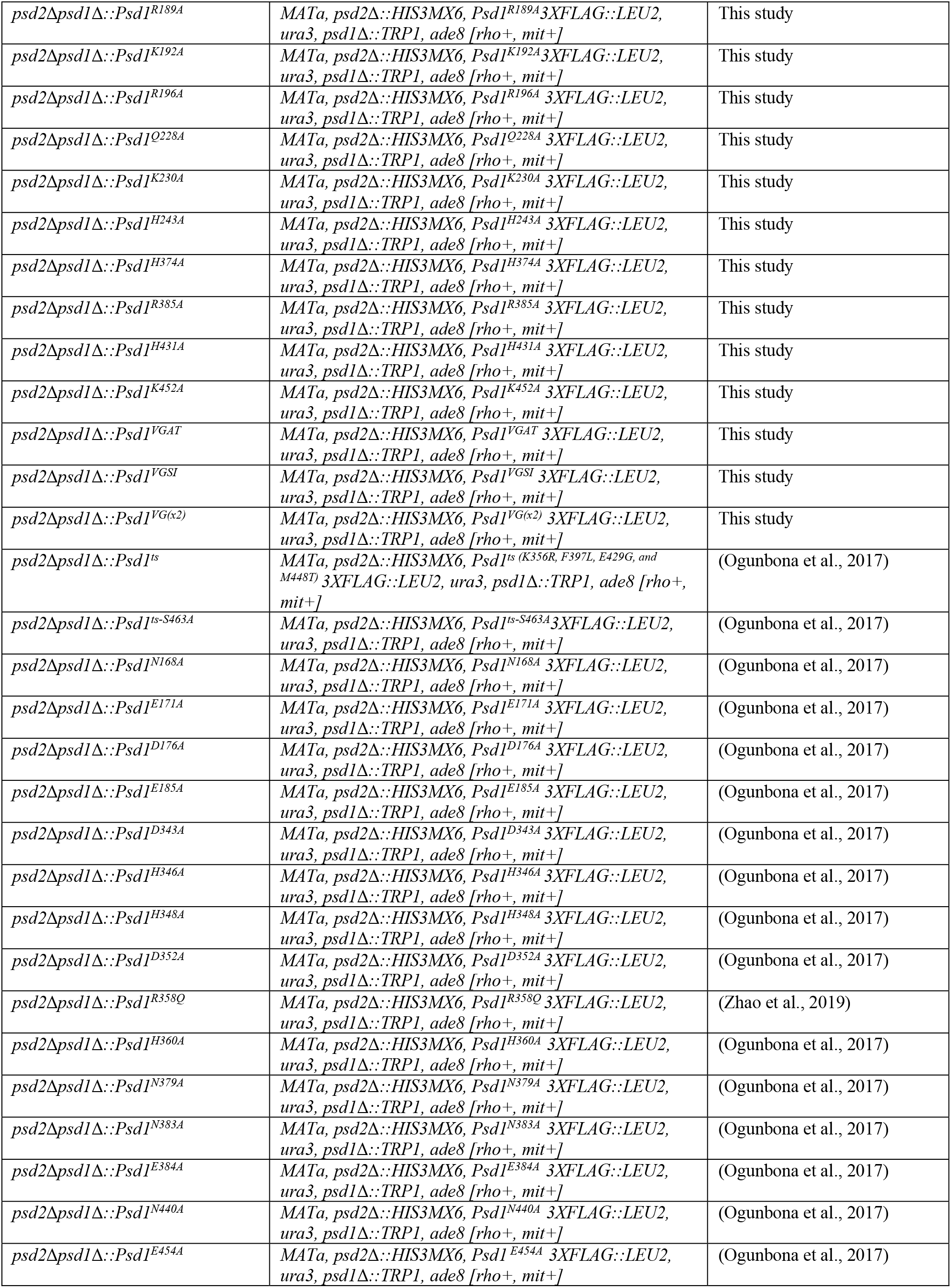
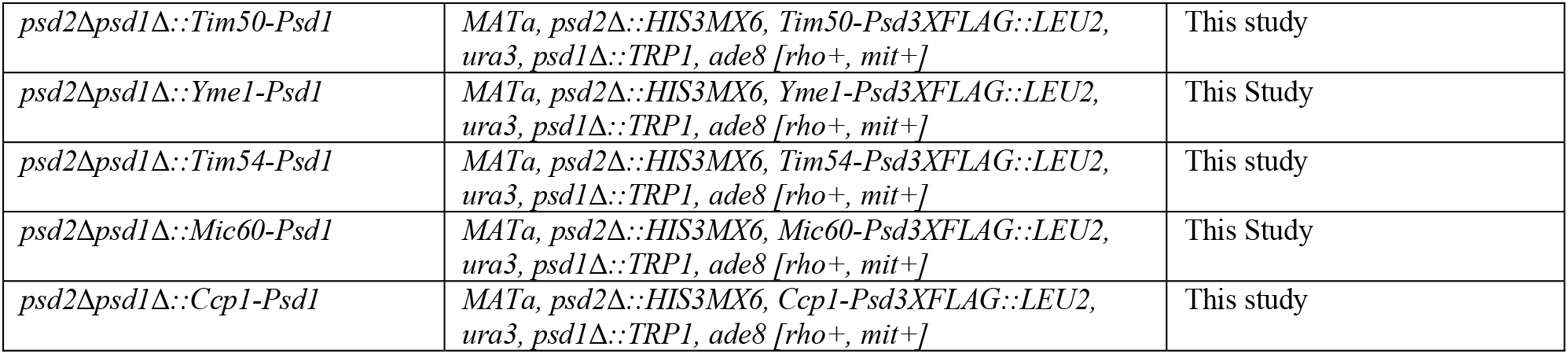
Yeast Strains.

Two approaches were used to generate deletion strains: 1) PCR-mediated gene replacement of the entire reading frame (Wach et al., 1994) and 2) homology-integrated (HI)-Clustered Regularly Interspaced Short Palindromic Repeats (CRISPR) gene editing in yeast, as previously described (Bao et al., 2015; Ogunbona et al., 2017). The *tam41*Δ strain and *pdr5*Δ strains were generated by replacing the entire *TAM41* or *PDR5* open reading frames with HISMX6 or CloNat/nourseothricin, respectively. All gene deletions were confirmed by either immunoblot (*tam41*Δ; Figure 7C) or genomic PCR (*pdr5*Δ). The *psd1*Δ*psd2*Δ*cho1*Δ strains was generated using a CRISPR-Cas9 gene block designed to target *CHO1*, as previously described (Calzada et al., 2019).

LGS/AAA mutations were inserted into the *PSD1* open reading frame of WT and *psd2*Δ yeast using a HI-CRISPR system as previously described (Bao et al., 2015; Calzada et al., 2019; Ogunbona et al., 2017). Briefly, the CRISPR-Cas9 gene block (gBlock) was designed using Benchling online CRISPR guide design tools to target *PSD1* and assembled into the plasmid pCRCT, a gift from Huimin Zhao (Addgene plasmid 60621), using the Golden Gate assembly method (Engler et al., 2009). The gBlock was composed of the CRISPR-Cas9 target (20bp), a 100 base pair (bp) homology-directed repair template with 50bp homology arms on either side flanking the Cas9 cutting site, and the codons encoding the LGS motif mutated to AAA by designing homology repair templates encoding these changes *psd1^LGS/AAA^* (<u>BsaI restriction site</u>; *homology arms*; **LGS/AAA mutations**; <u>*3’ CRISPR target*</u>; 5’-CTTT<u>GGTCTC</u>ACCAAAAC*TATTGGGAGGGATGCCTTTGGTTAAGGGTGAAGAAATGGGT GGCTTTGAA***GCGGCAGCC***ACTGTTGTACTTTGTTTTGAAGCTCCCACTGAATTTAAGTTCGATGTTAG<u>GAAATGGGTGGCTTTGAATT</u>*GTTTTAGAGA<u>GAGACC</u>TTTC-3’). The gblock was assembled into the pCRCT plasmid using the Golden Gate assembly method (Engler et al., 2009), and successfully confirmed by DNA sequencing. Following transformation of targeting plasmid into WT or *psd2*Δ yeast, positive clones were identified by immunoblot.

To determine if the various IM-directed Psd1 chimeras were functional, yeast strains were grown overnight in YPD and equal amounts (serial 1:4 dilutions starting at 0.00012 OD_600_’s in 3μl of sterile water) spotted onto SCD plates in the absence or presence of 2mM ethanolamine hydrochloride. To assess respiratory capacity, serially diluted strains were spotted on synthetic complete ethanol glycerol (SCEG; 0.17% yeast nitrogen base, 0.5% ammonium sulfate, 0.2% (w/v) complete amino acid mixture, 1% (v/v) ethanol, 3% (v/v) glycerol), and synthetic complete lactate (SC-LAC; 0.17% yeast nitrogen base, 0.5% ammonium sulfate, 0.2% (w/v) complete amino acid mixture, 0.05% (w/v) dextrose, 2% (v/v) lactic acid, 3.4mM CaCl2-2H2O, 8.5mM NaCl, 2.95mM MgCl2-6H2O, 7.35mM KH2PO4, and 18.7mM NH4Cl, pH 5.5) plates. Plates, which contained 2% (w/v) agar (Himedia, cat RM301), were incubated at 30°C or 37°C for the indicated days.

For routine growth, yeast were grown in synthetic complete dextrose medium (SCD; 0.17% yeast nitrogen base, 0.5% ammonium sulfate, 0.2% (w/v) complete amino acid mixture, 2% (w/v) dextrose) with or without 2mM ethanolamine, or YPD (1% (w/v) yeast extract, 2% (w/v) tryptone, 2% (w/v) dextrose) unless otherwise noted. For growth assays involving various supplements, overnight pre-cultures of the indicated yeast strains grown in YPD were inoculated to an OD_600_ of 0.2 in YPD media alone, or YPD supplemented with 2mM ethanolamine, 2mM choline, 1% (v/v) Tergitol (Type NP-40), 1% (v/v) Tergitol + 0.5mM lyso-PE, or 10 μM CCCP, and grown at 30°C overnight with vigorous shaking.

### Quantitation of mitochondrial Psd1

Quantification of Psd1 levels in three different batches of mitochondria isolated from WT or *psd1*Δ yeast grown in rich lactate medium (1% (w/v) yeast extract, 2% (w/v) tryptone, 0.05% (w/v) dextrose, 2% (v/v) lactic acid, 3.4mM CaCl2-2H2O, 8.5mM NaCl, 2.95mM MgCl2-6H2O, 7.35mM KH2PO4, 18.7mM NH4Cl, pH 5.5) was determined by immunoblot using a recHis6Psd1, generated and purified previously (Tamura et al., 2012a), standard curve.

### Preparation of cell lysates for immunoblot analysis

Unless otherwise indicated, the equivalent of 2.0 OD_600_ of yeast cells were harvested by centrifugation at 1690 *x g* for 10 min. Cell pellets were resuspended in 1mL of MilliQ grade water and treated with a final concentration of 120mM sodium hydroxide (NaOH) and 1% (v/v) β-mercaptoethanol (β-ME) ice with periodic inversion for 10 min on. 100% (w/v) trichloroacetic acid (TCA) was added for a final concentration of 6% and proteins precipitated on ice for 10 min with periodic inversion. Pellets were collected by centrifugation at 21,000 *x g* for 2 min, washed with 1 mL 100% acetone, and re-centrifuged at 21,000 *x g* for 2 min. Pellets were dissolved with 60μL 0.1M NaOH at room temperature prior to addition of 60μL 2X reducing sample buffer (RSB) and denaturation at 95°C for 5 min. Equal sample volumes were resolved by SDS-PAGE, transferred to nitrocellulose membranes, and analyzed by immunoblot.

### EndoH treatment in whole cells

Yeast cultures maintained on YPD plates were grown overnight in SCD or YPD liquid medium prior to harvesting 10 OD_600_ of each culture by centrifugation at 1690 *x g* for 10 min. Cell pellets were resuspended in 250μL of lysis buffer (20mM HEPES-KOH pH 7.4, 150mM potassium acetate, 2mM magnesium acetate, 1mM phenylmethylsulfonyl fluoride (PMSF)) and transferred to a 1.5mL microfuge tube containing ~50μL of glass beads. Samples were vortexed at level 10 for 30 min at 4°C. Unbroken cells were separated by centrifugation at 425 *x g* for 5 min and the supernatant transferred to a microfuge tube containing 10μL of 10% (w/v) sodium dodecyl sulfate (SDS) and 2μL of β-ME (final concentration 0.5% (w/v) SDS and 1% (v/v) β-ME). Samples were incubated at 95°C for 10 min and cooled at room temperature. Each sample was divided into two 100μL aliquots and diluted with an equal volume of 0.5M sodium citrate pH 5.5. One aliquot served as an untreated control; 1μL of Endoglycosidase H (EndoH; New England BioLabs) was added to the other sample. Samples were then incubated for 4 hrs at 37°C. Proteins were precipitated from the entire sample by addition of 100% (w/v) TCA to a final concentration of 20% and incubated on ice for 10 min. Pellets were collected by centrifugation at 21,000 *x g* for 10 min at 4°C and resuspended in 40μL of 0.1M NaOH and 40μL of 2X RSB as described above. Equal volumes of untreated and EndoH-treated samples were resolved by SDS-PAGE, transferred to nitrocellulose membranes, and analyzed by immunoblot.

### Mitochondrial isolation and fractionation

Isolation of crude mitochondria and cell fractionation by gravity centrifugation was performed as previously described (Claypool et al., 2006). Briefly, strains were cultured in 150mL of designated media shaking at 30°C for ~36 hours, then 100 OD_600_ was transferred to two Fernbach flasks containing 950mL of the same medium used for pre-cultures which were then grown at 30°C overnight for an OD_600_ of ∼2.5–3.5. Cell pellets, which were collected by centrifugation for 5min at 6,000 *x g*, were resuspended in water and re-collected at 2000 *x g* for 5min. The washed yeast pellet was resuspended in 50mL of 0.1M Tris-SO4, pH 9.4 containing 15mM dithiothreitol, and incubated for 20min at 30°C shaking at 220rpm. Following a 5 min spin at 2000 *x g*, yeast pellets were washed with 40mL of 1.2M sorbitol, 20mM KPi, pH 7.4, collected again at 2000×*g* for 5min, and resuspended with 1.2M sorbitol, 20mM KPi, pH 7.4 (2 mL/gram of yeast) spiked with Zymolyase 20T (3 mg/gram of yeast; Nacalai Tesque, Inc.) and shaken at 220rpm for 1h at 30°C. Following centrifugation at 3500 *x g* for 5min at 4°C, pellets were washed with 1.2M sorbitol, 20mM KPi, pH 7.4 and collected at 3500×*g* for 5min at 4°C; this washing step was performed twice. The final washed pellet was resuspended in 50mL of 0.6M sorbitol, 20mM KOH-MES, pH 6.0 (BB6.0 buffer) containing 1mM phenylmethylsulfonyl fluoride (PMSF) and subjected to 15 strokes in a tight-fitting (type A) glass dounce kept on ice. 200μL was set aside and analyzed as the starting material (SM) for cell fractionation studies. Cell homogenates were centrifuged for 5min at 1700 *x g* and the supernatant set aside. The pellet was again resuspended with ~50 mL BB6.0 containing PMSF, subjected to 15 strokes in a tight-fitting glass dounce kept on ice, and centrifuged for 5min at 1700 *x g*. The resulting supernatant was combined with that generated from the first round of dounces. The combined supernatants were centrifuged for 10min at 13,500 *x g* to sediment mitochondria-enriched pellets (P13). The P13 fraction was resuspended in 35mL of BB6.0, homogenized with two strokes of a telfon pestle, spun at 1700 *x g* 5min to remove debris from the supernatant which was again centrifuged at 13,500 *x g* for 10min. The same procedure was performed a second time with the resulting pellet, except that 0.6M sorbitol, 20mM HEPES-KOH, pH 7.4 (BB7.4) was used in place of BB6.0 and PMSF was omitted. When additional subcellular fractions were collected, 35mL of the supernatants from the original 13,500 *x g* centrifugation step was centrifuged at 21,500 *x g* for 15min to remove remaining mitochondria (P21), and the resulting supernatant centrifuged at 40,000 *x g* for 30min to separate ER-enriched membranes (P40) from light membranes and cytosol (S40). Protein concentration was determined using the Pierce BCA Protein Assay Kit (ThermoFisher Scientific, Catalog No. 23225) and aliquots of SM, P13, P21, P40 and S40 were frozen in liquid nitrogen and stored at −80°C.

### EndoH Treatment in crude mitochondria and microsomal fractions

100μg of each P13 and P40 sample was resuspended with 20μL of 1X glycoprotein denaturing buffer (New England BioLabs) and incubated at 100°C for 10 min. For removal of N-glycans, samples were incubated in a final volume of 40μL containing 1X G5 reaction buffer (New England BioLabs) and incubated at 37°C for 4 hrs in the absence (mock) or presence of 1μL EndoH. 30μg of each sample was resolved by SDS-PAGE and analyzed by immunoblot.

### TMT Proteomics

#### Protein processing

P40 samples, collected through subcellular fractionation and quantified by BCA assay as described above, were resuspended and denatured in 8 M urea with 100 mM ammonium bicarbonate, pH 7.8. Disulfide bonds were reduced by incubation for 45 min at 57°C with a final concentration of 10 mM Tris (2-carboxyethyl) phosphine hydrochloride (Catalog no C4706, Sigma Aldrich). A final concentration of 20 mM iodoacetamide (Catalog no I6125, Sigma Aldrich) was then added to alkylate these side chains and the reaction was allowed to proceed for one hour in the dark at 21°C. Aliquots of 100 μg protein were taken and diluted to 1 M urea using 100 mM ammonium bicarbonate, pH 7.8. Trypsin (V5113, Promega) was added at a 1:100 ratio and the samples were digested for 14 hours at 37 °C.

#### Tandem Mass Tag (TMT) labeling and cation exchange-based fractionation of peptides

The digested lysates were desalted using C18 Omix tips (Cat number A57003100, Agilent) and dried down in 50 μg portions. For each of the conditions, 50 μg of digested peptides were labeled with the TMT 10plex reagent (Cat number 90113, ThermoFisher). Dried pellets were resuspended in 50 μL of 1X phosphate buffered saline and to each tube, one aliquot of the respective TMT reagent, which was resuspended in 41 μL of acetonitrile, was added. The reaction was allowed to proceed at 21°C for one hour and then quenched via the addition of 8 μL of 1 M ammonium bicarbonate. The samples were dried down and then desalted using Omix tips. The individual TMT labeled samples were pooled and then fractionated using strong cation exchange chromatography on an AKTA Pure 10 (GE Healthcare) equipped with a Luna 5 μm 100 angstrom 150 x 2.1 mm strong cation exchange (SCX) column (Catalog no 00F-4398-B0, Phenomenex). Buffer A was 5 mM KH2PO4 in 30% acetonitrile (Catalog no 34998, Sigma Aldrich), pH 2.7. Buffer B was 350 mM KCl (Catalog no PX1405 EM Science) in buffer A. A 200 μL/min gradient was run from 0% B to 50% B over 10 mL, then up to 100% B over 1 mL. A total of 12 fractions were collected during the peptide elution portion of the gradient.

#### Mass spectrometry (MS)

Individual fractions from the strong cation exchange (SCX) chromatography were desalted using ZipTips (ZTC18S096, EMD Millipore), dried down and resuspended in 0.1% formic acid (Catalog no 94138, Honeywell). Fractions were analyzed by LC-MS on an Orbitrap Fusion Lumos (ThermoFisher) equipped with an Easy NanoLC1200 HPLC (ThermoFisher). Peptides were separated on a 75 μm × 15 cm Acclaim PepMap100 separating column (Thermo Scientific) downstream of a 2 cm guard column (Thermo Scientific). Buffer A was 0.1% formic acid in water. Buffer B was 0.1% formic acid in 80% acetonitrile. Peptides were separated on a two-hour gradient from 0% B to 35% B. Peptides were collisionally fragmented using HCD mode. Precursor ions were measured in the Orbitrap with a resolution of 120,000. Fragment ions were measured in the Orbitrap with a resolution of 50,000. The spray voltage was set at 2.2 kV. Orbitrap MS1 spectra (AGC 1×106) were acquired from 400-1800 m/z followed by data-dependent HCD MS/MS (collision energy 42%, isolation window of 0.7 Da) for a three second cycle time. Charge state screening was enabled to reject unassigned and singly charged ions. A dynamic exclusion time of 60 seconds was used to discriminate against previously selected ions.

#### Database search

The LC-MS/MS data was searched against a *Saccharomyes cerevisiae* (strain ATCC 204508 / S288c) database downloaded from Uniprot on 4/30/2018. Proteome Discoverer version 2.1.1.21 (ThermoScientific) was used to interpret and quantify the relative amounts from the diagnostic fragment ions of the TMT reagent. The database search parameters were set as follows: two missed protease cleavage sites were allowed for trypsin digested with 5 ppm precursor mass tolerance and 0.02 Da for-fragment ion quantification tolerance. Trypsin was set as the protease with up to two missed cleavages allowed. Oxidation of methionine, pyroglutamine on peptide amino termini and protein N-terminal acetylation were set as variable modifications. Carbamidomethylation (C; +57Da) was set as a static modification. TMT 10plex was set as a constant modification on peptide amino termini and lysine residue side chains. Data was searched using the Sequest HT algorithm and the results were filtered via Percolator with a decoy database false discovery rate (FDR) set to < 1% as a filter for peptide identification (Spivak et al., 2009). Volcano plots were generated by RStudio Version 1.3.959.

### Overexpression of PISD in HEK cells

HEK cells were cultured in DMEM (Gibco) with 10% heat-inactivated FBS (Gibco). Cells were seeded at 2 × 10^6^ cells per plate on 10-cm plates 24 hours prior to transfection. Cells were transfected using 30 μL of Lipofectamine 3000 (Thermo Fisher Scientific) reagent and 30 μL of P3000™ Reagent with 15 μg of plasmid DNA per plate. Media was changed 48 hours after transfection. Cells were collected for immunofluorescence 72 hours after transfection.

### Immunofluorescence

Cover slips were fixed in phosphate-buffered saline buffer containing 4% (v/v) paraformaldehyde at 37°C for 15 min followed by quenching with ammonium chloride (50 μM). Fixed cover slips were permeabilized in 0.25% (v/v) Triton X-100. Samples were incubated with the following primary antibodies: anti-FLAG (NB600-344; Novus Biological) for PISD, anti-TOMM20 (Sigma; HPA011562) for detection of mitochondria, and anti-calnexin (Millipore; MAB3126) for detection of ER. Appropriate Alexa Fluor–labelled secondary antibodies (Invitrogen) were applied next. Finally, cover slips were mounted for imaging with Dako fluorescent mounting medium (S3023; Agilent Technology).

### Imaging and image analysis

Images were captured using 488 nm, 568 nm and 640 lasers on Olympus spinning disc confocal system (Olympus SD-OSR) using 100x lens (UAPON 100XOTIRF) running MetaMorph software in super-resolution mode. Images were originally captured as 11 z-stacks spanning 0.2 μm between each stack. Image stacks were compressed to a single image using max intensity projection and combined into a composite image using ImageJ. To obtain the intensity plot, a line selection was made in the composite image, pixel intensity profile was obtained along the line selection using “get profile” function for each channel and plotted in a line graph.

### Phospholipid Analyses

To determine the steady phospholipid profiles in *psd2*Δ*psd1*Δ transformed with the assorted IM-directed Psd1 chimeras, starter cultures were diluted to an OD_600_ = 0.4 in 2 mL of rich lactate medium supplemented with 10 μCi/mL ^32^Pi and grown shaking at 240 rpm for ∼24 hr in a 30°C water bath. For the remaining phospholipid analyses, starter cultures were diluted to an OD_600_ = 0.1 in 2 mL of YPD supplemented with 0.5 μCi/mL ^14^C-Acetate and grown shaking at 240 rpm for ∼24 hr in a 30°C water bath. Where indicated, cultures additionally contained 2 mM choline, 2mM ethanolamine or 1% (v/v) Tergitol + 0.5mM lyso-PE. Yeast were centrifuged at 1690 *x g* for 5 min, washed with 2 mL MilliQ grade water and re-centrifuged at 1690 *x g* for 5 min. The yeast pellets were resuspended in 0.3 mL MTE buffer (0.65M mannitol, 20mM Tris, pH 8.0, and 1 mM EDTA) containing 1mM PMSF, 10 μM leupeptin, and 2 μM pepstatin A, transferred to a 1.5 mL microcentrifuge tube containing 0.1 mL glass beads, and each tube sealed with parafilm. Yeast were disrupted by vortexing on high for ∼30 min at 4 °C. Glass beads and unbroken yeast were removed after a 2 min 250 *× g* centrifugation at 4°C. The resulting extract was used to assess cellular phospholipid levels. To sediment a crude mitochondrial pellet, the cellular extract was centrifuged for 5 min at 13,000 *× g* at 4°C. Following liquid scintillation, equal amounts of labeled cell extract or crude mitochondria were transferred to 5 mL borosilicate tubes containing 1 mL chloroform and 0.5 mL methanol and vortexed for 30 min on medium-high at room temperature. Phase separation was initiated by adding 0.3 mL 0.9% (w/v) NaCl to each sample which was then vortexed on high for 1 min. Samples were centrifuged at 1000 rpm in a clinical centrifuge for 5 min at room temperature, and the upper aqueous phase aspirated into radioactive waste. The organic phase was washed with 0.25 mL 1:1 Methanol:H2O, vortexed on high for 30 seconds at room temperature, and centrifuged as before. The lower organic phase was transferred to a new tube and dried down under a stream of nitrogen. Just prior to resolving the phospholipids by thin layer chromatography, dried lipid extracts were resuspended in 13 μL of chloroform and loaded on SILGUR-25 (Machery-Nagel) TLC plates that had been pretreated with 1.8% (w/v) boric acid in 100% ethanol and activated at 95 °C for at least 30 min. Phospholipids were resolved using chloroform/ethanol/H2O/triethylamine (30:35:7:35) after which the plates were air-dried for 30 min and developed using a K-screen and FX-Imager (Bio-Rad Laboratories).

### Reverse transcription-quantitative real-time PCR

Yeast were grown in YPD at 30°C to an OD_600_ of 0.5-0.7, and RNA was extracted using hot phenol extraction as previously described (Amin-ul Mannan et al., 2009). 1uL of Turbo DNase (TURBO DNA-free Kit; Invitrogen) was used to remove DNA from 10 μg of RNA, RNeasy MiniElute Cleanup Kit (Qiagen) to purify RNA, and SuperScript VILO Master Mix (Invitrogen) to reverse-transcribe 1 μg of RNA into cDNA. The qPCR was done exactly as described in (Ogunbona et al., 2018) using the FastStart Universal SYBR Green Master Rox (Roche), a 1:10 dilution of synthesized cDNA as template, and 100nM of primers for a total reaction volume of 20 μL. The primers used are *CIS1* forward, 5’-AATGCAGGAGTTGCTTACA-3’; *CIS1* reverse, 5’-TTCTTCCGCTATAGTCTCCAA-3’; *PSD1* forward, 5′-CCAGTAGCACAAGGCGAAGA-3′; *PSD1* reverse, 5′-GACATCAAGGGGTGGGAGTG-3′; and the housekeeping genes *ACT1* forward, 5′-GTATGTGTAAAGCCGGTTTTG-3′; and *ACT1* reverse, 5′-CATGATACCTTGGTGTCTTGG-3′. For controls, we included no-template and minus-RT controls. Samples were run using the QuantStudio 6 Flex Real-Time PCR System (Thermo Fisher). The Ct value differences between *ACT1* and target genes (*PSD1* or *CIS1*) were computed to compare mRNA levels in the tested strains.

### Submitochondrial localization

The submitochondrial localization assay as previously described (Claypool et al., 2006), determines the accessibility of proteins to protease (100μg/mL Proteinase K) in intact mitochondria, mitoplasts with osmotically ruptured OMs, and detergent solubilized mitochondria. For intact mitochondria, 150 μg of mitochondria were placed in two tubes and resuspended in 1mL of BB7.4 lacking or containing Proteinase K. In parallel, 600μg of mitochondria was centrifuged at 8,000 *x g* for 5min at 4°C and the resulting pellet resuspended in 200μL of BB7.4 and then aliquoted into four microfuge tubes, each containing 50μL. To rupture the OM only, 950 μL of 20mMK^+^HEPES pH 7.4 lacking or containing Proteinase K was added to two tubes which were vortexed on medium for 15 sec. To provide access to all mitochondrial compartments, 950 μL of 0.5% (w/v) deoxycholate in 20mMK^+^HEPES pH 7.4 lacking or containing Proteinase K was added to the final two tubes which were vortexed on medium for 15 sec. All samples were incubated on ice for 30 minutes after which 20 mM PMSF was added prior to centrifuging each sample at 21,000×*g* for 10min at 4°C. For intact mitochondria and OM-ruptured mitoplasts, after aspirating the supernatant, the pellet was resuspended in 180μL of BB7.4 + 1mM PMSF and transferred to a new tube that contained 20μL of 100% (w/v) TCA. For detergent solubilized mitochondria, the supernatants were transferred to new tubes that contained 0.2mL of 100% (w/v) TCA. All tubes were incubated on ice for 1h, and centrifuged at 21,000 *x g* for 10min at 4°C. After the resulting supernatants were discarded, the pellets were washed with 0.5mL of cold acetone, spun at 21,000×*g* at 4°C for 2 minutes, and the supernatants aspirated. 30μL of 0.1M NaOH was added to tubes to resuspend the pellets, after which 30 μL of 2X RSB was added. Samples were incubated at 95°C for 5 minutes and then resolved by SDS-PAGE and analyzed by immunoblot.

### Aggregation Assay

The detergent based aggregation assay was performed as done previously (PMID: 21300850). In brief, 200 μg of mitochondria was centrifuged at 21,000 *x g*, 4°C for 5 min to pellet mitochondria. Post-aspirating the supernatant, the pellets were resuspended in 40 μL 1.5% (w/v) digitonin lysis buffer (20 mM HEPE-KOH, pH 7.4, 100 mM NaCl, 20 mM Imidazole, 1mM CaCl2, 10% glycerol) containing 1mM PMSF, 10 μM leupeptin, and 2 μM pepstatin A. The samples were incubated on ice and flicked to mix every 10 min for 30 min. Next, samples were centrifuged at 21,000 *x g*, 4°C for 10 minutes and the supernatants (S1) transferred to new tubes containing 50μL 2X RSB. Pellets were resuspended with 40 μL 1% (v/v) TX-100 lysis buffer (20 mM HEPE-KOH, pH 7.4, 100 mM NaCl, 20 mM Imidazole, 1mM CaCl2, 10% glycerol) supplemented with 1mM PMSF, 10 μM leupeptin, and 2 μM pepstatin A, kept on ice and flicked to mix every 10 min. After 30 min, samples were centrifuged at 21,000 *x g*, 4°C for 10 minutes and the supernatants (S2) transferred to new tubes containing 50μL 2X RSB; the pellet (P2) was resuspended in 90 μL of 1X RSB. All samples were incubated for 5 min at 95°C, resolved by SDS-PAGE and analyzed by immunoblot.

### Homology model

Homology modeling of Psd1 based on the *E. coli* PSD structure (PDB code: 6L06) was performed using SWISS-MODEL (https://swissmodel.expasy.org) (Waterhouse et al., 2018). Psd1 sequence (residues 120-500), except the signal sequence and the transmembrane region at the N-terminus, was used for modeling.

### Cycloheximide and proteasome inhibition experiments

Cycloheximide (CHX) degradation experiments were performed essentially as previously described (Claypool et al., 2011). Starter cultures of yeast grown in YPD at 30°C overnight were diluted to an OD_600_ = 1 in a total volume of 10 mL YPD or YPD + 2 mM choline and incubated for 5 min at 30°C shaking at 220 rpm. To inhibit cytosolic protein synthesis, CHX was added at 200 μg/mL. For the MG132 drug treatment, starter cultures of yeast grown in YPD at 30°C overnight were collected and then resuspended to an OD_600_ = 1 in a total volume of 10 mL SCD + 2mM choline. Following the addition of 80 μM of MG132 or an equivalent volume of DMSO, samples were incubated for 1 hour at 30°C shaking at 220 rpm prior to the addition of 200 μg/ml of CHX. At the indicated time points, 1 OD_600_ units of cells were transferred to a tube containing an equal volume of ice cold 2X azide mixture (20 mM NaN3 and 0.5 mg/ml bovine serum albumin) and yeast pellets collected following 1690 *x g* centrifugation for 10 min at 4°C. The pellets were frozen at −80°C until all time points were collected at which point cell lysates were harvested as already described.

### RNA-Sequence

Yeast strains were grown in YPD (D), YPD containing 2 mM ethanolamine (E), YPD containing 2 mM choline (C), or YPD containing 1% (v/v) tergitol and 0.5 mM lyso-phosphatidylethanolamine (L), with shaking at 220 rpm, to an OD_600_ between 0.5-0.7. RNA was extracted using the hot phenol extraction as previously described (Amin-ul Mannan et al., 2009). At least 50 ng/μl (for a total of 2 μg) of RNA per sample were sent to Novogene for quality tests, library construction, sequencing, and data analysis, detailed below.

#### Data analysis

*Saccharomyces cerevisiae* release 98 reference genome and gene model annotation files were downloaded from genome website browser (NCBI/UCSC/Ensembl) directly. Indexes of the reference genome was built using STAR and paired-end clean reads were aligned to the reference genome using STAR (v2.5). HTSeq v0.6.1 was used to count the read numbers mapped of each gene. FPKM of each gene was calculated based on the length of the gene and reads count mapped to this gene. Differential expression analysis between two conditions/groups (three or four biological replicates per condition) was performed using the DESeq2 R package (2_1.6.3). The resulting p-values were adjusted using the Benjamini and Hochberg’s approach for controlling the False Discovery Rate (FDR).

#### Heatmaps

Genes presented in Figures 7E and 7F exhibited the largest degree of increase or decrease in fold change (FC) in LGS mutant (∷LGS) YPD versus WT (∷WT) YPD, all having adjusted p-values of less than 0.05. Genes encoding retrotransposons, putative proteins of unknown function, or those corresponding to dubious open reading frames were excluded. Heatmaps were created with Prism 8.

#### Data access

The high-throughput sequencing data from this study have been submitted to the Gene Expression Omnibus under accession number GSE162987.

### *In Vivo* Retro-translocation assay

The *in vivo* retro-translocation assay was adapted and modified from (Neal et al., 2018). Cells were grown to log phase (OD_600_ 0.2-0.3) at 30°C and 15 OD_600_’s of cells were pelleted. Cells were resuspended in H20, centrifuged and lysed with the addition of 0.5 mm diameter silica beads (Biospec) and 400μL of XL buffer (1.2 M sorbitol, 5 mM EDTA, 0.1 M KH2PO4, final pH 7.5) with protease inhibitors, followed by vortex in 1 min intervals for 6-8 min at 4°C. Lysates were combined and clarified by centrifugation at 2,500 *x* g for 5 min. Clarified lysate was ultracentrifuged at 100,000 *x* g for 15 min to separate pellet (P100) and supernatant fraction (S100). P100 pellet was resuspended in 200μL of 1.5% (w/v) digitonin lysis buffer (20 mM Tris-Cl, pH 7.4, 100 mM NaCl, 20 mM Imidazole, 1mM CaCl2, 10% glycerol) with protease inhibitors and 5 mM N-ethyl maleimide (NEM, Sigma) and incubated for 2 hours at 4°C followed by centrifugation of extract for 30 min at 4°C. Clarified extract was transferred to 600 μL of basic lysis buffer (20mM HEPES-KOH, pH7.4, 100mM NaCl, 20mM Imidazole, 1mM CaCl2, 10% glycerol) with protease inhibitors and NEM containing 30 μL 1:1 anti FLAG Affinity Gel (Genscript). S100 supernatant was added directly to 600 μL of basic lysis buffer with protease inhibitors and NEM containing 30 μL 1:1 anti FLAG Affinity Gel (Genscript). All samples were incubated overnight at 4°C. Samples were washed once with 0.1% (w/v) Digitonin wash buffer (20 mM Tris-Cl, pH 7.4, 100 mM NaCl, 20 mM Imidazole, 1mM CaCl2), washed once with 0.1% (w/v) High Salt Digitonin wash buffer (20 mM Tris-Cl, pH 7.4, 250 mM NaCl, 20 mM Imidazole, 1mM CaCl2) and washed once more with 0.1% (w/v) Digitonin Low Salt wash buffer (20 mM Tris-Cl, pH 7.4, 20 mM Imidazole, 1mM CaCl2), aspirated to dryness, and resuspended in 2X Urea sample buffer (8 M urea, 4% SDS, 1mM DTT, 125 mM Tris, pH 6.8), and incubated at 55°C for 10 min. Samples were resolved by 8% SDS-PAGE, transferred to nitrocellulose, and immunoblotted with monoclonal anti-ubiquitin (Fred Hutchinson Cancer Center, Seattle), anti-FLAG for Psd1-α, anti-Psd1(Tamura et al., 2012a) for Psd1-β, Mito β-α, and ER β-α, and goat anti-rabbit or anti-mouse (Bio-Rad) conjugated with horseradish peroxidase (HRP) to recognize the primary antibodies. Western Lightning^®^ Plus (Perkin Elmer, Watham, MA) chemiluminescence reagents were used for immunodetection.

### Proteolytic removal of ubiquitin from retrotranslocated Psd1 LGS mutant

Ubiquitin removal was accomplished with the broadly active Usp2 ubiquitin protease as previously described (Garza et al., 2009), except that human recombinant Usp2Core (LifeSensors Inc., Malvern, PA) was used, and leupeptin and NEM were excluded from all buffers. Briefly, 100 μL of S100 supernatant containing the *in vivo* retrotranslocated Psd1-LGS mutant was incubated with 20 μL of Usp2Core (5 μg) for 1 hr at 37°C. The reaction was quenched with 200 μL of SUME (1% SDS, 8 M Urea, 10 mM MOPS, pH 6.8, 10 mM EDTA) with protease inhibitors and retrotranslocated Psd1-LGS mutant was immunoprecipitated as described above. 20 μL of the bound material was used for detection of Psd1-LGS mutant with anti*-*Psd1 antibody.

### Antibodies

Antibodies employed in this study include rabbit antibodies specific to yeast Psd1 (Tamura et al., 2012a), Qcr6 (Baile et al., 2013), Hsp70 (Claypool et al., 2006), Pic1 (Whited et al., 2013), Tom70 (Baile et al., 2013), Tim54 (Hwang et al., 2007), Abf2 (Calzada et al., 2019), Kgd1 (Glick et al., 1992), Cho1 (Choi et al., 2010), Kar2 (Huyer et al., 2004); and mouse monoclonal antibodies against Sec62 (gift of David Meyers, University of California, Los Angeles, Los Angeles, CA), Aac2 (Panneels et al., 2003), FLAG (clone M2, catalog number F3165, Sigma), Ubiquitin (clone P4D1, catalog number sc-8017, Santa Cruz Biotechnology), and Cox2 (Anti-MTCO2 antibody [4B12A5] ab110271, Abcam). Mature Tam41 (Arg35-stop codon) was cloned into pET28a (Novagen) downstream of the encoded His6 tag, induced in BL21-CodonPlus(DE3)-RIL *Escherichia coli* and affinity purified with Ni^2+^ agarose (Qiagen). The purified His6Tam41 protein was used as an antigen in rabbits (Pacific Immunology). Other antibodies used were horseradish peroxidase-conjugated (Thermo Fisher Scientific), and fluorescence-conjugated secondary antibodies (Pierce).

### Miscellaneous

Quantity One Software (Bio-Rad Laboratories) was used to quantify immunoblots and TLC plates. Statistical comparisons were performed using SigmaPlot 11 software (Systat Software, San Jose, CA) or Prism 8 (GraphPad); *P* values of ≤0.05 were deemed significant. All the graphs show means, and error bars are standard errors of the mean (SEM). All figure panels are representative of at least three biological replicates, unless otherwise notes.

## ACKNOWLEDGEMENTS

We would like to thank Drs. Jonathan Friedman (UT Southwestern, USA) and Jodi Nunnari (UC Davis, USA) for sharing W303-JF WT and Δ*psd1* strains, Drs. Susan Michaelis (JHMI, USA), George Carman (Rutgers University, USA), and Carla Koehler (UCLA, USA) for antibodies, Dr. Hilla Weidberg (University of British Columbia, Canada) for sharing her protocol for detecting *CIS1* transcript levels, and Linhao Ruan and Drs. Selvaraju Kandasamy, Oluwaseun B. Ogunbona, Eric Spear, and Samuel Jayakanthan for technical assistance. This work was supported by the National Institutes of Health (Grant R01GM111548 to S.M.C.), the National Science Foundation Graduate Research Fellowship Program (DGE1746891 to P.N.S.), a Biochemistry, Cellular, and Molecular Biology Program training grant (T32GM007445 to E.C.), a predoctoral fellowship from the American Heart Association (16PRE31140006 to M.G.A.), the Indiana University Precision Health Grand Challenge Initiative to J.C.T.; NIH grant (1R35GM133565 to S.E.N) and Pew Biomedical Award (to S.E.N.), the Japan Society for the Promotion of Science (JSPS) KAKENHI (20K15734 to Y.W.), and the Canadian Institutes of Health Research (T.E.S.).

## Author Contributions

S.M.C conceived the project; P.N.S., E.C., T.Z., A.N., T.S., S.E.N., and S.M.C. designed research, performed experiments and made strains; P.N.S., M.G.A., and S.M.C. analyzed data; J.C.T performed the mass spectrometry-based proteomics; Y.W. developed and analyzed the homology models of Psd1; and P.N.S., E.C., and S.M.C. wrote the paper with the feedback and approval from all authors.

---

**Figure S1.**
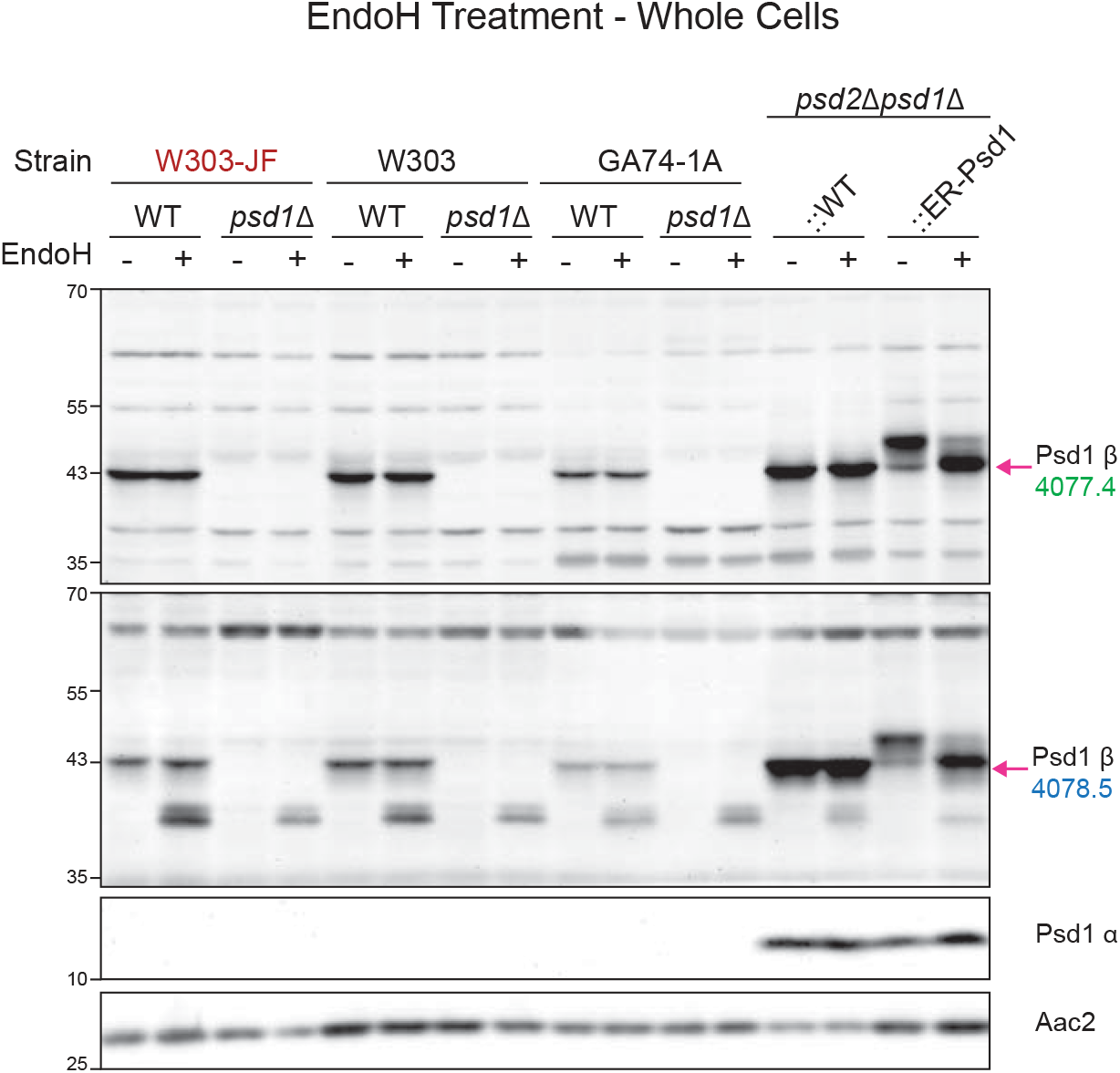
Absence of glycosylated Psd1 in strains used by Friedman et al. Cell extracts derived from pairs of WT and *psd1*Δ yeast of the indicated backgrounds grown at 30°C in synthetic complete dextrose (SCD) were treated with EndoH as listed and analyzed by immunoblot using the designated anti-Psd1 antisera; Aac2 served as loading control. W303-JF is a kind gift from Jonathan Friedman (n=3).

**Figure S2.**
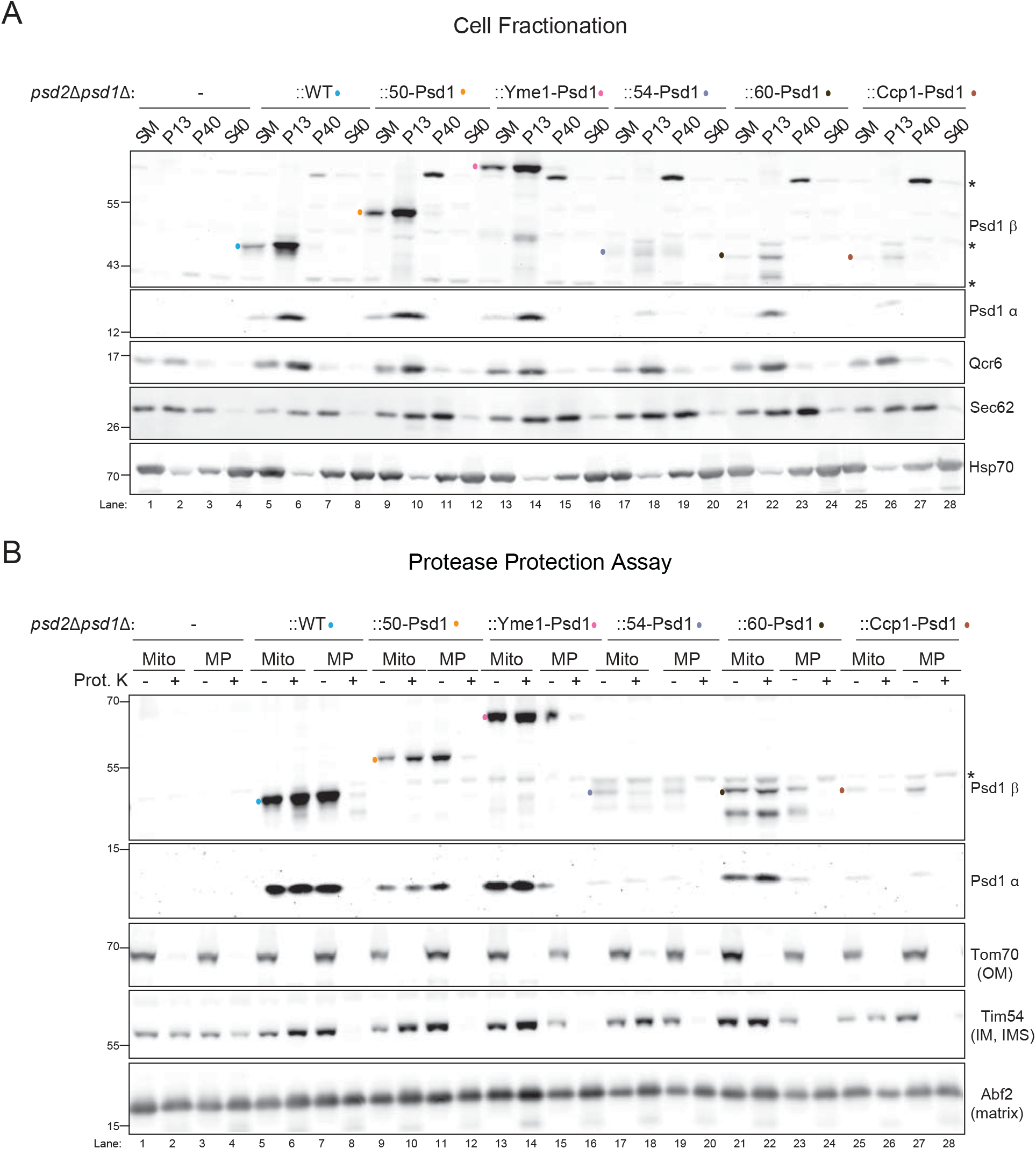
Assorted IM-Psd1 chimeras are properly localized. (A) Subcellular fractions were collected from the indicated strains following growth in YPD medium. Equal amounts of each fraction were resolved by SDS-PAGE and immunoblotted for both subunits of Psd1 (β and α) mitochondrial (Qcr6), ER (Sec62), and cytosolic (Hsp70) markers. SM, starting material, P13, pellet of 13,000xg; P40, pellet of 40,000xg; and S40, supernatant of 40,000xg (n=3). (B) Intact mitochondria (Mito) and osmotically ruptured mitochondria (MP) from indicated yeast strains were treated as indicated with 100 μg proteinase K (Prot. K) and the resultant samples resolved by SDS-PAGE and immunoblotted for Psd1 (β and α subunits) and the compartment-specific markers Tom70 (OM), Tim54 (IMS), and Abf2 (matrix). For Psd1 immunoblots, different protein amounts, 10 μg (lanes 5-16) or 40 μg (lanes 1-4 and 17-28), were resolved due to differences in chimera abundance (n=3). *, nonspecific bands.

**Figure S3.**
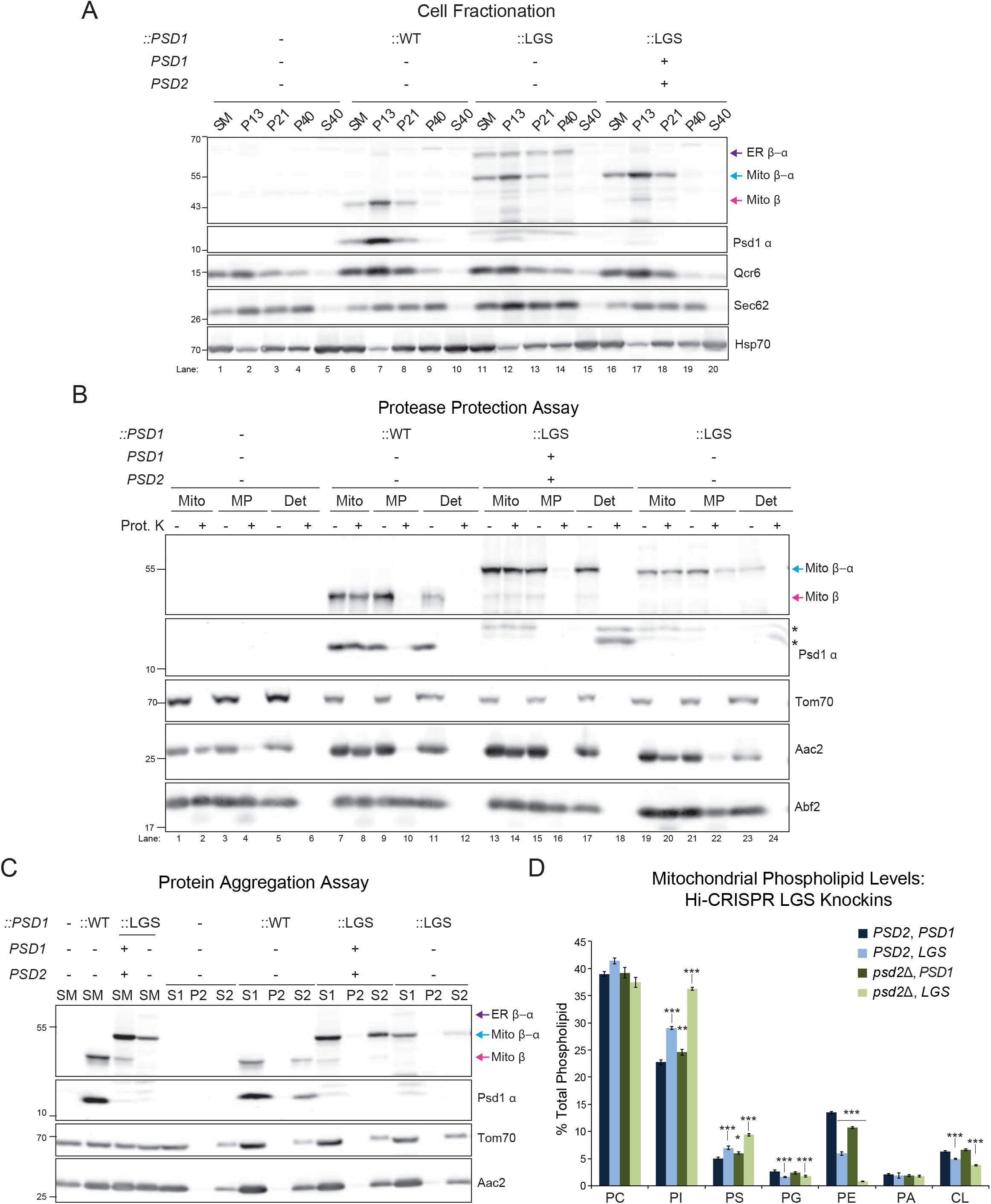
Glycosylated mutant Psd1 co-fractionates with the ER and is not aggregation prone. (A) Subcellular fractions were collected from the indicated strains following growth in YPD medium. Equal amounts of each fraction were resolved by SDS-PAGE and immunoblotted for both subunits of Psd1 (β and α), and mitochondrial (Qcr6), ER (Sec62), and cytosolic (Hsp70) markers. SM, starting material, P13, pellet of 13,000x*g*; P40, pellet of 40,000x*g*; and S40, supernatant of 40,000x*g* (n=3). (B) Protease protection assay in intact mitochondria (Mito), osmotically ruptured mitochondria (MP), or deoxycholate-solubilized mitochondria (Det.) treated as indicated with 100 μg proteinase K (Prot. K). The resultant samples were resolved by SDS-PAGE and immunoblotted for Psd1 (β and α subunits), and the compartment-specific markers Tom70 (OM), Aac2 (IMS), and Abf2 (matrix). Aac2 was detected using a monoclonal antibody, 6H8, which recognizes an N-terminal epitope present in the IMS (n=3). *, nonspecific bands. (C) Mitochondria isolated from the indicated strains grown in YPD at 30oC were solubilized with digitonin and separated into a supernatant (S1) and pellet (P1) by centrifugation. The resulting pellet (P1) was re-extracted with TX-100 and again fractionated into a supernatant (S2) and pellet (P2) by centrifugation. Collected fractions were resolved by SDS-PAGE and immunoblotted as indicated. SM, starting material (n=3). (D) Mitochondrial phospholipid levels from control and CRISPR-LGS knockin strains analyzed in Fig. 3G (mean ± SEM for n=6). Significant differences (1 symbol P ≤ 0.05; 2 symbols P ≤ 0.01; 3 symbols P ≤ 0.001) compared to WT (*PSD2*, *PSD1*) were calculated by one-way ANOVA with Holm-Sidak pairwise comparison.

**Figure S4.**
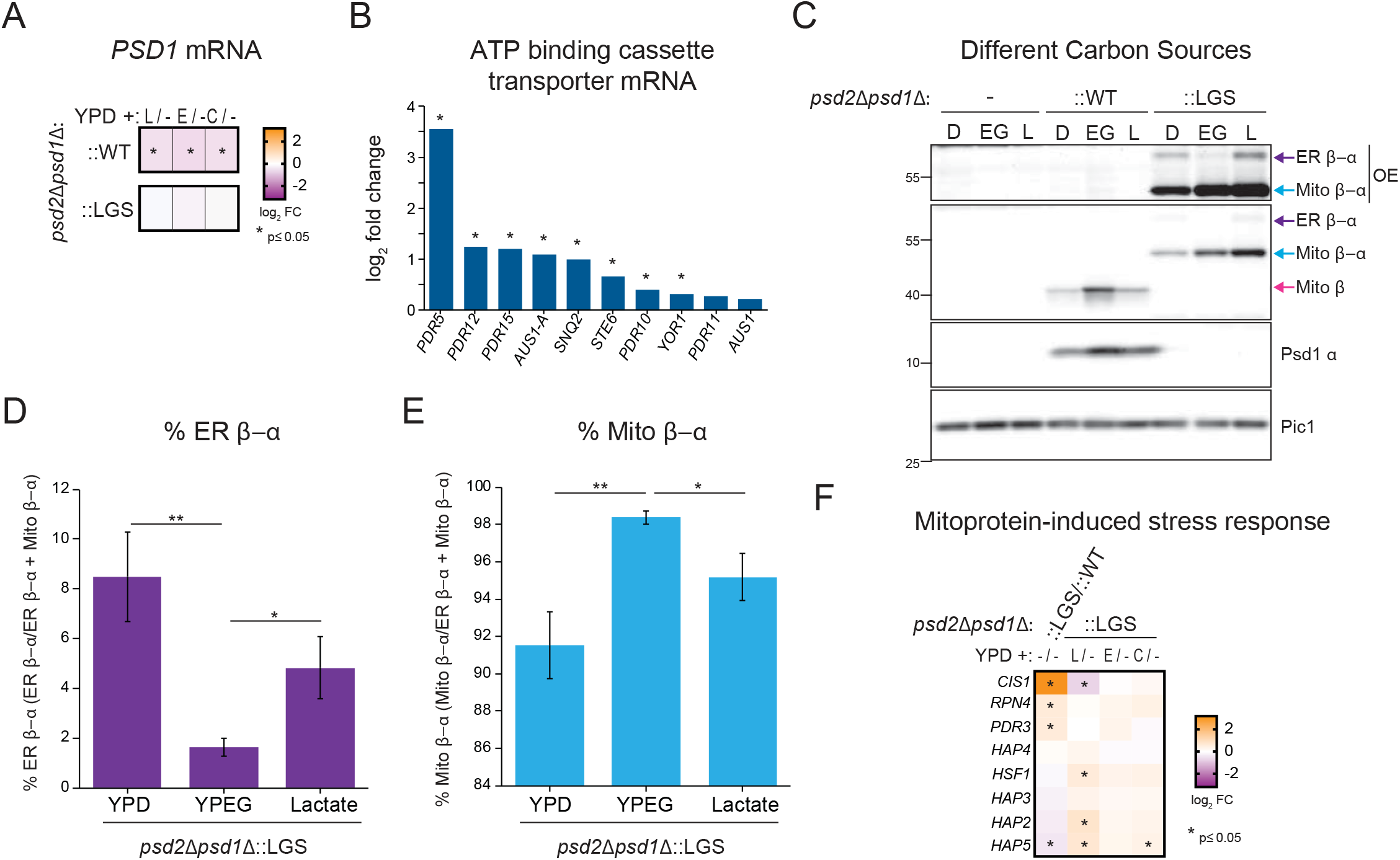
Additional transcriptomic and metabolic comparisons. (A) *PSD1* mRNA levels in ∷WT and ∷LGS yeast cultured in dextrose alone compared to dextrose supplemented with LPE (L), ethanolamine (E), or choline (C). padj values that were significant (P ≤ 0.05) are designated (*). (B) Relative abundance of ATP binding cassette transporter genes in ∷LGS vs ∷WT (*symbols padj values P ≤ 0.05). (C) The indicated strains were pre-cultured at 30°C in YPD (D), YPEG (EG) and Rich Lactate (L) and after isolation of cell extracts, the α and β subunits of Psd1 were analyzed by immunoblotting. Pic1 served as a loading control (n=4). OE, over-exposed. (D and E) The relative abundance of (D) ER β-α, and (E), Mito β-α in Psd1-LGS expressing yeast was determined (means ± SEM for n=4). Statistical differences (1 symbols P ≤ 0.05; 2 symbols P ≤ 0.01) were determined by unpaired Student’s t-test. (F) Heatmaps of Mitoprotein-induced stress genes in ∷LGS vs ∷WT and how each supplement does or does not affect their expression relative to ∷LGS grown in YPD alone. padj values that were significant (P ≤ 0.05) are designated (*).

